# Aberrant hippocampal transmission and behavior in mice with a stargazin mutation linked to intellectual disability

**DOI:** 10.1101/2021.06.08.447333

**Authors:** GL Caldeira, AS Inácio, N Beltrão, CAV Barreto, MV Rodrigues, T Rondão, R Macedo, RP Gouveia, M Edfawy, J Guedes, B Cruz, SR Louros, IS Moreira, J Peça, AL Carvalho

## Abstract

Mutations linked to neurodevelopmental disorders, such as intellectual disability (ID), are frequently found in genes that encode for proteins of the excitatory synapse. Transmembrane AMPA receptor regulatory proteins (TARPs) are AMPA receptor auxiliary proteins that regulate crucial aspects of receptor function. Here, we investigate an ID-associated mutant form of the TARP family member stargazin. Molecular dynamics analyses showed that the stargazin V143L variant weakens the overall interface of the AMPAR:stargazin complex and hinders the stability of the complex. Knock-in mice for the V143L stargazin mutation manifest cognitive and social deficits and hippocampal synaptic transmission defects. In the hippocampus of stargazin V143L mice, CA1 neurons show impaired spine maturation in basal dendrites, and synaptic ultrastructural alterations. These data demonstrate a causal role for mutated stargazin in the pathogenesis of ID and highlight its role in the development and function of hippocampal synapses.

## INTRODUCTION

Most of the fast component of excitatory neurotransmission in the central nervous system is mediated by glutamate receptors of the α-amino-3-hydroxyl-5-methyl-4-isoxazole-propionate type (AMPAR). This family of receptors is associated with receptor auxiliary proteins that regulate their traffic, gating and pharmacology, thus increasing receptor functional diversity in the brain ^1-3^. Members of the transmembrane AMPAR regulatory protein (TARP) family are widely expressed AMPAR auxiliary subunits ^4^, and key modulators of AMPAR-mediated transmission. The prototypical TARP stargazin (also known as TARP γ2) was discovered in the ataxic stargazer mouse ^5^, which lacks synaptic AMPARs on cerebellar granule cells ^6^. Stargazin interacts with both AMPA receptor subunits and synaptic PDZ-containing proteins such as postsynaptic density protein 95 [PSD95; ^7^], and this is required for targeting AMPAR to synapses ^7,8^. The stargazin/PSD95 complex has recently been found to form a condensed assembly via liquid-liquid phase separation, a process that is critical for AMPAR-mediated transmission ^9^.

TARPs, including stargazin, couple with the majority of AMPAR complexes in the brain, promote receptor trafficking to the cell surface and their synaptic targeting, and augment their functional properties ^1,2^. Stargazin slows the rates of AMPAR desensitization and deactivation thus increasing the size of the postsynaptic current. Not surprisingly, stargazin regulates baseline synaptic transmission and is also involved in Hebbian and homeostatic forms of synaptic plasticity that are dependent on tightly regulated AMPAR traffic ^10-12^.

Impaired glutamatergic synaptic transmission and plasticity have been implicated in neurodevelopmental disorders ^13^. Evidence from human genetic studies suggests that copy number variation or the presence of rare point mutations in genes encoding proteins of the ionotropic glutamate receptor complex may play a role in the aetiology of these disorders ^14-18^. Single nucleotide polymorphisms in the *CACNG2* gene encoding stargazin were associated with a subgroup of schizophrenia patients ^19^, and alterations in the DNA copy number and in the levels of stargazin mRNA were detected in the post-mortem brain of schizophrenia patients ^20,21^. Dysregulated stargazin expression was also found in the dorsolateral prefrontal cortex of patients with bipolar disorder ^21^, and stargazin polymorphisms were associated with the response to lithium, a frequent treatment for bipolar disorder ^21,22^. A *de novo* missense mutation in *CACNG2* has been identified in a non-syndromic intellectual disability (ID) patient with moderate severity ^16^. Taken together, these data point to a possible link between stargazin and the pathogenesis of neurodevelopmental disorders, which has not yet been investigated. Evaluating how human mutations in the stargazin-encoding gene disrupt synaptic function and impact behavior may also provide insight into the physiological role of stargazin.

Here, we investigated the impact of the ID-associated missense V143L mutation in stargazin ^16^ in the molecular dynamics (MD) of the AMPAR:stargazin complex, in the cell surface diffusion of stargazin and in its ability to traffic AMPAR to the neuronal surface and to the synapse. To evaluate behavioral, neuronal morphology and functional alterations triggered by the stargazin V143L variant, we generated a knock-in (KI) mouse model to express the mutant protein. We found that stargazin V143L KI mice display alterations in cognitive and social behavior, along with altered hippocampal spine morphology, associated with synaptic ultrastructural defects. We also found disrupted synaptic transmission and aberrant stargazin phosphorylation in stargazin V143L mutant mice.

## RESULTS

### Intellectual disability-associated stargazin V143L mutation affects the AMPAR:stargazin complex structure

A *de novo* missense mutation in the *CACNG2* gene encoding stargazin was described in a heterozygous 8 year-old male patient with moderate, non-syndromic, intellectual disability ^16^. This mutation leads to substitution of valine143 by leucine (p.V143L), a residue in the third transmembrane domain of stargazin that is highly conserved among species (Figs. 1a,c), suggesting a critical role for the function of stargazin. Accordingly, the V143L substitution was predicted to be damaging using PolyPhen-2 ^23^, SIFT ^24^ and PROVEAN ^25^. Importantly, the stargazin V143L variant has not been described in databases collecting sequencing variants for the general population (Genome Aggregation Database or Exome Variant Server).

**Figure 1.**
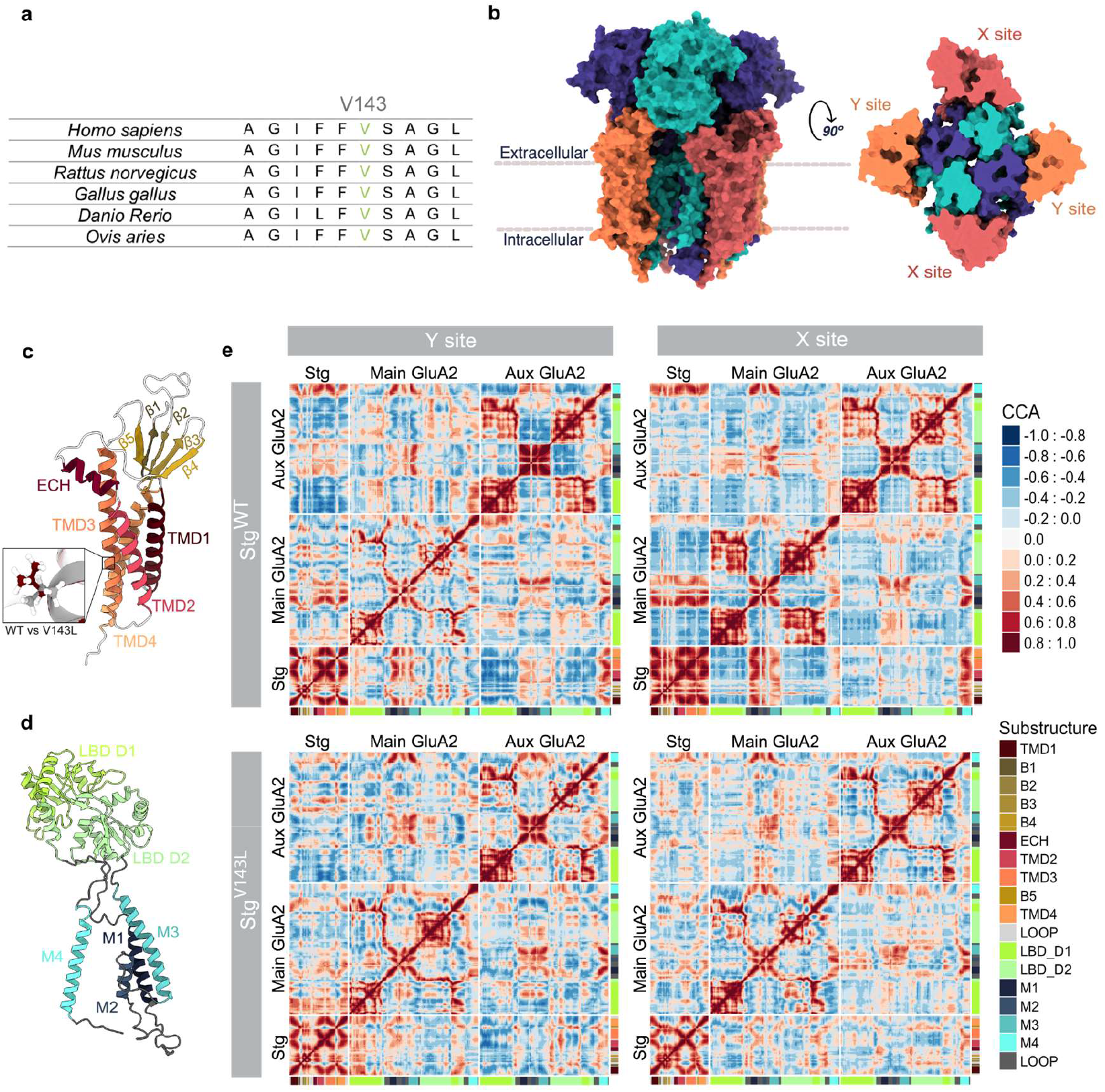
The ID-associated stargazin mutation in the highly conserved V143 residue weakens the interaction between stargazin and AMPARs. **(a)** Valine 143 in stargazin is highly conserved among species. **(b)** Surface representation of AMPAR:stargazin complex viewed parallel to the membrane (left) and from the extracellular side (right). The extracellular view of the complex (at the membrane level) shows the two different sets of stargazin assembly points (X and Y sites) around AMPAR. Each GluA subunit is colored individually in shades of blue. Stargazin molecules are colored in orange (Y site) or brick (X site). **(c)** Side view of the stargazin structure shown as a cartoon with substructures labelled and colored in a spectrum of yellow/orange. Close-up shows the V143L mutation (WT – grey; ID – red). **(d)** Side view of a GluA2 subunit structure shown as cartoon with substructures labelled and transmembrane domains colored in a spectrum of blue and ligand-binding domain colored in green. **(e)** Dynamical cross-correlation maps for the AMPAR:stargazin complex WT form in Y site and X site and for stargazin V143L form in the Y and X sites. Substructure annotation was added at the bottom and right of each map for easier reading. CCA goes from −1 (anticorrelated, opposite direction) to 1 (correlated, same direction). See also Figs. S1-S4.

In order to characterize the effect of the V143L stargazin mutation in the structure and dynamics of the AMPAR:stargazin complex, we used molecular dynamics (MD) simulations. MD simulations are key to attain an atomistic understanding of biomolecular processes from ligand-binding to protein-coupling induced conformational changes, and are widely used to interpret experimental data or/and to guide experimental work ^26,27^. Nevertheless, to the best of our knowledge, possibly due to the high number of residues involved, there are no MD studies available regarding AMPAR:TARP complexes. To perform the first MD study on this complex, we took advantage of increasing availability of structural data, along with more readily accessible computational resources, to apply MD algorithms and predict *in silico* how the AMPAR:stargazin system responds to a particular perturbation. To this purpose we used homology modeling (Fig. 1b) to construct both the WT and V143L models of the AMPAR:stargazin complex, based on one of the described cryo-EM structures for the complex ^28^.

AMPARs are composed of variable multidomain subunits (GluA1-GluA4, Fig. 1d) arranged in a “Y” shape, divided into three layers: the amino-terminal domains (ATDs), the ligand-binding domains (LBDs), and the transmembrane domain (TMD). TMDs, with four helices (M1-M4), show a pseudo-fourfold symmetric structure, a two-fold rotational symmetry formed by two dimers composed of A/D and B/C subunits, providing high conformational flexibility ^29,30^. As such, AMPAR:TARPs shows variable stoichiometry with an apparent maximum of four TARPs that can be broken down into two groups: TARPs binding X sites (common interfaces with AMPAR subunits A/B or C/D) and Y sites (involving subunits A/D or B/C) [^30^, Fig. 1b]. We will herein refer to the GluA structure exhibiting the highest contact surface with the coupled TARP as *Main GluA* and *Auxiliary GluA* to the other one. We analyzed the effect of the stargazin V143L mutation at both sites taking into account different metrics on macromolecular rearrangements such as cross-correlation analysis (CCA) and root mean square deviations (RMSD), and also at the interfacial level (solvent accessible surface area -SASA).

Figure 1e reveals the network of correlated/anticorrelated (same/opposite direction) motions between different regions of the the AMPAR:stargazin complex structure, which informs on the impact of the stargazin V143L mutation in the overall structural conformation of the macromolecular complex. The dynamics between X and Y sites are distinct. The loops between TMD1-TMD2 and TMD3-TMD4 in stargazin are highly anticorrelated with the rest of the structure, whereas in the X site the movements are widely positively correlated. In the *Main GluA*, LBD-D1 and LBD-D2 are anticorrelated for the Y site and positively correlated for the X site. In the stargazin V143L system these differences are attenuated. In the complex containing stargazin V143L, both sites are much closer to the Y site’s motions of the complex containing the WT protein. Correlation analysis for the complete system shows a positive correlation between M1-M2-M3 of *Main GluA* and M4 of the *Auxiliary GluA* with the transmembrane regions of stargazin in both X and Y sites. This correlation weakens and, in some regions, shifts to an anticorrelation with the introduction of the V143L stargazin mutation, especially in the M2-M3 of *Main GluA* and M4 of *Auxiliary GluA* (observed in both sites). In the Y site, the β1-β4 region in stargazin displays a stronger positive correlation with the D2 of the *Main GluA* when compared to the X site, which is also weakened in the mutated system. Finally, in the Y site, stargazin has a slight positive correlation with the *Auxiliary GluA* transmembrane domain that is lost in the mutated systems. In the X site this correlation is never present. Overall CCA results indicate that the X site is more prone to conformational rearrangements introduced by the V143L mutation in stargazin than the Y site.

RMSD was used to assess protein conformational changes between different time points in the trajectory, and their distribution is shown in Figures S1 and S2 for all monomers at both sites, in the WT and mutated complexes. Comparing WT-and V143L stargazin-containing mutated complexes, the maximum RMSD values do not change significantly, but their density is lower for lower values in the complex containing stargazin V143L, demonstrating a higher flexibility. RMSD values for stargazin main substructures, α-helices and β-strands, are illustrated in Figures S2a and S2b, respectively. TMD3 and TMD4 show higher density for lower RMSD values, and therefore lower conformational flexibility in the mutated system. In the case of the β-strands, their RMSD values were higher for the mutated system at both sites, which shows a higher conformational flexibility of this system, especially for β4 and β5. For GluA, at both X and Y sites, the M1 and M2 show higher deviation and conformational flexibility in the mutated systems, while M3 and M4 show lower deviation and flexibility, particularly for the mutated X site.

The effect of the V143L stargazin mutation on the size of the coupling interface between stargazin and AMPAR was further assessed by measuring the sum of all residues SASA (Figs. S3 and S4). For both sites and systems, M1 in the *Main GluA* and M4 in the *Auxiliary GluA* display the lowest ΔSASA values (highest contribution) in the TM region, with M1 presenting a different behavior between the two sites (data not shown). In stargazin, the substructures with the highest contribution for the interface were TMD3 and TMD4 in the TM region (data not shown). When comparing average values per substructure in the WT and V143L mutant forms of stargazin, the LBD_D2 of *Auxiliary GluA*, β4 of *stargazin* and the loops regions show significant differences (Fig. S3). Analysis of ΔSASA per component of the interface (*Main GluA, Auxiliary GluA* and stargazin) shows significant dissimilarities for stargazin in the X site, and for *Main GluA* in the Y site (Fig. S4).

Taken together, the MD analysis indicates that the V143L mutation in stargazin weakens the interaction of stargazin with the AMPAR complex, particularly in the X interaction site.

### The V143L mutation affects the trafficking properties of stargazin

Stargazin plays a role in AMPAR trafficking through the early compartments of the biosynthetic pathway ^31^, and mediates complexed AMPAR trafficking to the cell membrane, their synaptic stabilization ^7,32^ and surface diffusion trapping ^8,33^ through binding to PSD95. Given the described roles, we explored the potential effect of the V143L mutation on stargazin’s cell surface diffusion properties. Low-density cortical neurons were co-transfected with plasmids encoding Homer-GFP, for synapse identification, and HA-tagged WT stargazin (StgWT), or the V143L stargazin variant (StgV143L). We monitored stargazin diffusion by single nanoparticle imaging of HA-stargazin using quantum dots (QDs) (Figs. 2a,b). Stargazin V143L particles displayed increased mean square displacement (MSD) (Fig. 2c), decreased synapse residence time (Fig. 2d) and higher global and synaptic diffusion coefficients than StgWT (Figs. 2e,f), suggesting that the V143L mutation renders stargazin more mobile in the plasma membrane.

**Figure 2.**
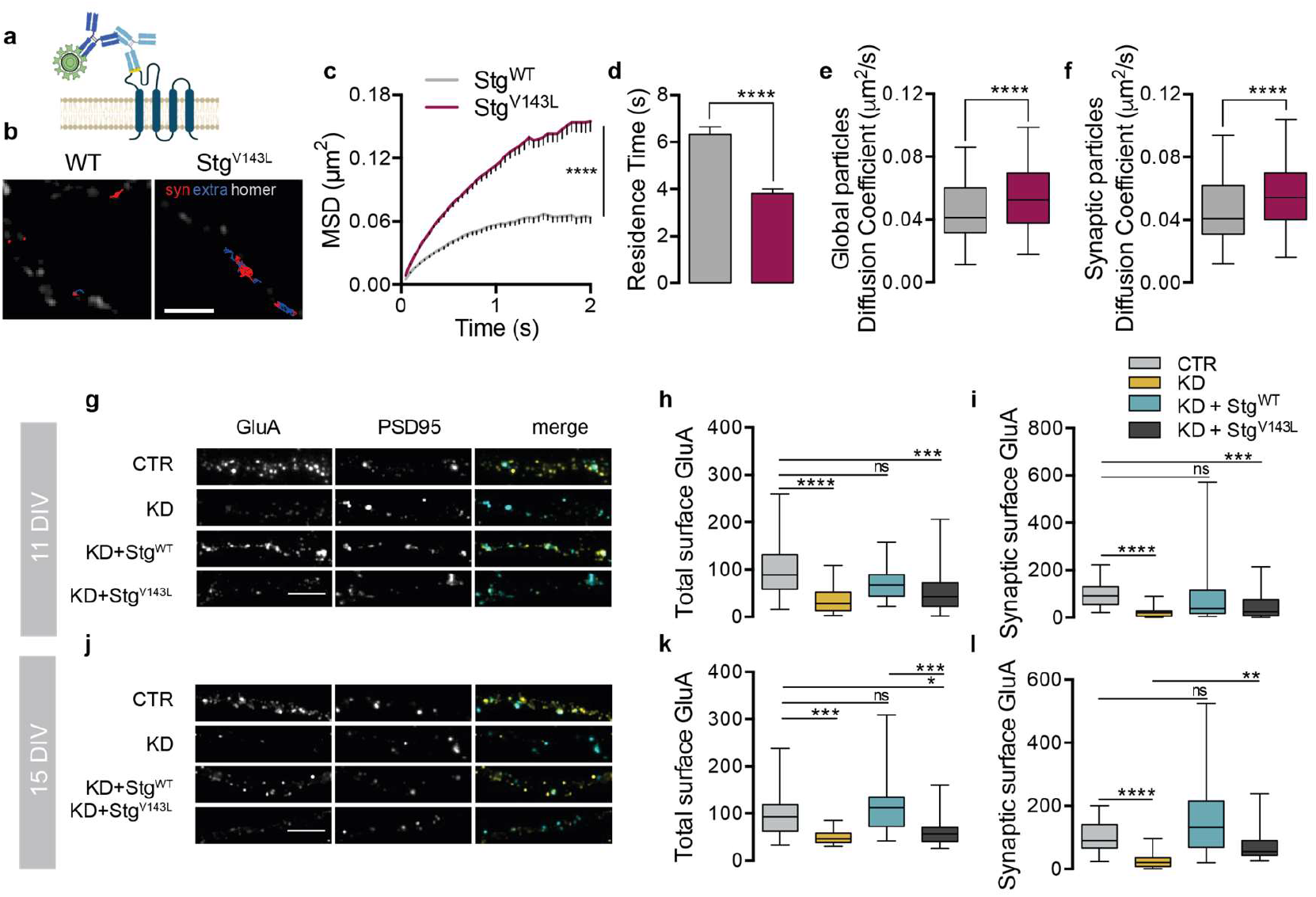
Stargazin ID-associated variant presents altered surface diffusion and elicits defective AMPAR trafficking. **(a)** Cortical neurons were co-transfected with Homer1C-GFP and HA-stargazin (either the WT form of stargazin – Stg^WT^ – or the intellectual disability-associated variant – Stg^V143L^). Stargazin surface diffusion was evaluated using quantum dot (QD)-labelled secondary antibodies (dark blue) against anti-HA antibodies (light blue) to detect the extracellular HA epitope (yellow) in stargazin (HA-stargazin). (**b**) Reconstructed HA-stargazin trajectories (synaptic and perisynaptic -red, extrasynaptic -blue) and Homer1C-GFP signal (white). Scale bar represents 5 µm. (**c**) Stargazin mean square displacement (MSD) (± SEM) versus time plots for cells expressing WT stargazin or the V143L variant. \*\*\*\**p* < 0.0001 by Mann Whitney test. (**d**) Mean synaptic residence time (± SEM) of WT stargazin and stargazin V143L. \*\*\*\**p* < 0.0001 by unpaired t-test. (**e, f**) Surface diffusion coefficient of global (**e**) and synaptic (**f**; Homer1C-GFP-colocalized) single QD-stargazin particles. Median diffusion (± 25%–75% IQR) of 277 and 485 for global trajectories, respectively and 139 and 280 for synaptic trajectories, respectively. \*\*\*\**p* < 0.0001 by unpaired t-test. A minimum of 18 cells were analyzed from 3 independent experiments. (**g-l**) Disrupted AMPAR surface expression in the presence of the stargazin V143L variant. Low-density cortical neurons were transfected at 7 (**g**,**h**,**i**) or 11 (**j**,**k**,**l**) days *in vitro* (DIV) with a control plasmid (pLL-shRNA-CTR) or with pLL-shRNA-Stg, which downregulates endogenous stargazin expression, or co-transfected with pLL-shRNA-Stg and pcDNA-Stg^WT^ or pcDNA-Stg^V143L^. Total surface and synaptic levels of GluA subunits were analyzed by immunocytochemistry at DIV11 or DIV15. (**g**,**j**) Representative images of GluA distribution and quantification of total (**h**,**k**) and synaptic (**i**,**l**) intensity of GluA clusters show impaired trafficking of AMPAR in neurons where stargazin was silenced, or which expressed the Stg^V143L^ variant. GluA accumulation at synaptic sites was assessed by the colocalization with PSD95 clusters. Clusters from 11 DIV cells were quantified from at least 33 cells imaged from four independent experiments; clusters from 15 DIV cells were quantified from 20 cells from two independent experiments. \*\*\*\**p* < 0.0001; \*\*\**p* <0.001; two-way ANOVA, followed by Dunn’s multiple comparison post hoc test. Boxes show the 25^th^ and 75^th^ percentiles, whiskers range from the minimum to the maximum values, and the horizontal line in each box shows the median value. Scale bars represent 5 μm.

The ID-associated mutation is located in the third transmembrane domain of stargazin (Fig. 1c), which was shown to be involved in the interaction with AMPAR subunits ^34,35^. Our molecular dynamics analyses indicate that this mutation weakens the interaction of stargazin with the AMPAR complex, in particular in the X site (Fig. 1e). We thus hypothesized that stargazin V143L may be defective in trafficking AMPAR to the cell surface and to the synapse. To test this possibility, we used a molecular replacement strategy in which we silenced endogenous stargazin expression in cultured cortical neurons and re-introduced either WT stargazin or the V143L variant. We assessed the effect of stargazin depletion and of the expression of the stargazin V143L variant in AMPAR trafficking and synaptic stabilization in young (Figs. 2g-i) and mature neurons (Figs. 2j-l). Low density rat cortical neurons were transfected with a control shRNA (CTR) or with a specific shRNA that depletes the levels of endogenous stargazin [KD; ^11^]. Cell surface and synaptic expression levels of AMPAR were evaluated by immunolabeling GluA AMPAR subunits using an antibody specific for their extracellular N-terminal region (Figs. 2g-l). As previously described ^11^, stargazin silencing led to a decrease on the cell surface (Figs. 2h,k) and synaptic levels of AMPAR (Figs. 2i,l). AMPAR clusters were considered synaptic when colocalizing with PSD95, whose expression was not affected by stargazin silencing (data not shown). In cells co-transfected with stargazin shRNA and WT shRNA-refractory stargazin (KD + StgWT), total and synaptic surface levels of GluA were rescued to basal levels. Critically, neuronal transfection of shRNA-refractory stargazin V143L mutant (KD + Stg^V143L^) led to a failure in mediating normal AMPAR traffic to the cell surface (Figs. 2h,k) and to the synapse (Figs. 2i,l), showing that the ID-associated mutation impairs stargazin’s role in AMPAR trafficking in both young and mature neurons.

### Genetically engineered mice with the stargazin V143L mutation show altered cognitive and social behavior

In order to study the effects of the ID-associated stargazin mutation *in vivo*, we generated a knock-in (KI) mouse line in which the human mutation was introduced in the mouse *Cacng2* gene. Using the gene targeting strategy we targeted the *Cacng2* gene to modify the nucleotide in the third exon which was found to be mutated in the ID patient ^16^. We designed a targeting vector containing two homology arms and a third segment containing the modification to be inserted in the mouse genome. In order to allow genotyping of the animals, a random sequence was inserted upstream the third exon (green sequence; Fig. 3a). Confirmation of the mutation was performed by Sanger sequencing (Fig. 3b). Heterozygous and homozygous KI mice were viable, did not display gross abnormalities, and did not show spontaneous seizures. To determine whether expression of the stargazin V143L variant affects gross brain morphology, we performed Nissl staining in brain coronal slices and compared sections from WT and homozygous stargazin V143L KI (KI^VL/VL^) mice. As shown in Figure 3c, no apparent macroscopic defects were visible in the brain of stargazin KI^VL/VL^ animals when compared with WT littermates, suggesting that overall brain morphology is not affected by expression of the stargazin V143L mutation. Moreover, the structural organization of the hippocampus and the cortical lamination were preserved (Figs. 3c and S5b,c).

**Figure 3.**
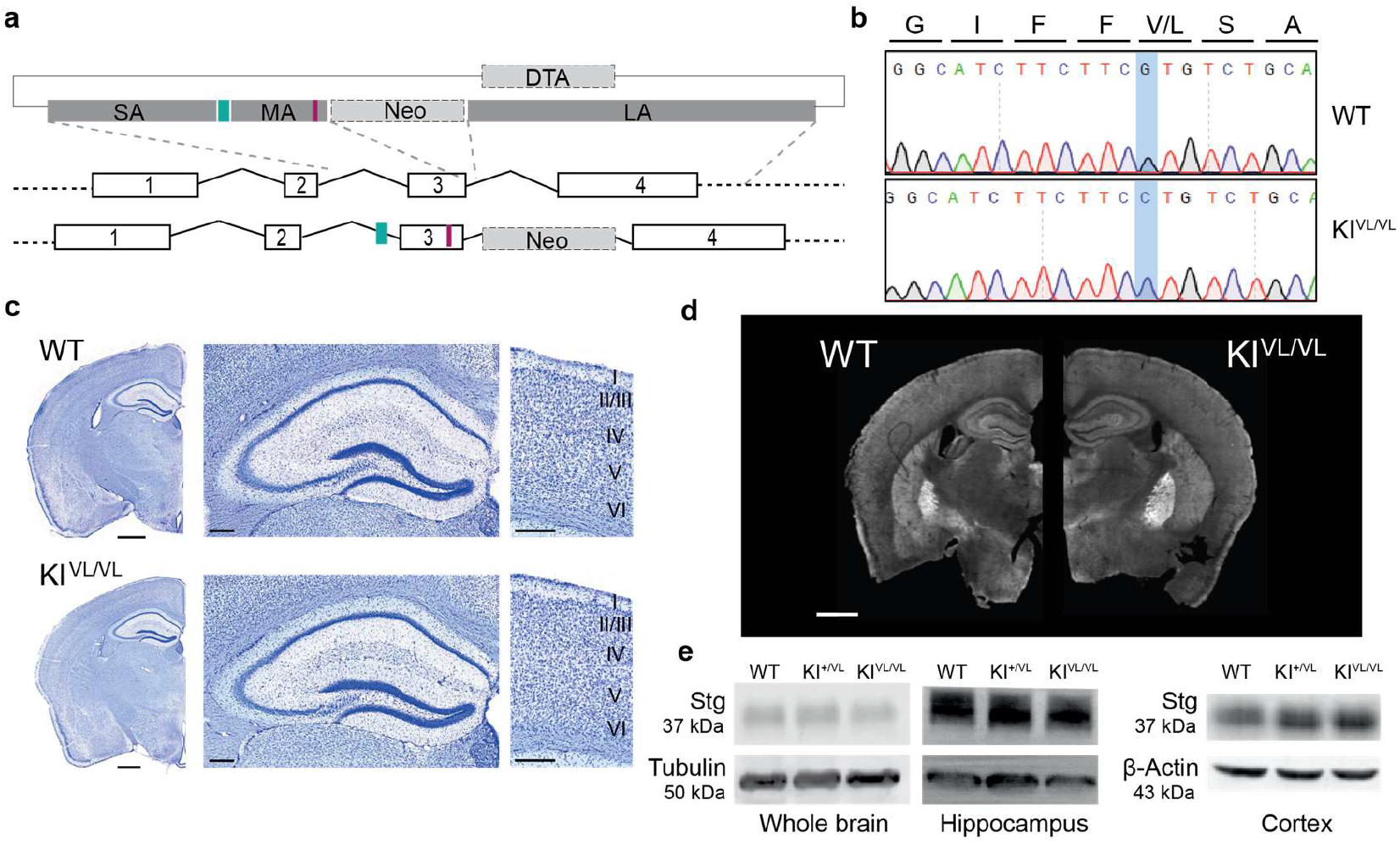
Stargazin V143L knock-in mice express normal stargazin levels and present no gross brain abnormalities. **(a)** Strategy for generating stargazin V143L knock-in mice. A vector containing two selection markers, Neo and DTA, and three homology arms was constructed: the short (SA) and the long arm (LA) allowed homologous recombination with the genomic DNA of mouse embryonic stem (ES) cells, and the middle arm (MA) contained the ID-associated point mutation (V143L) (red bar). A forward primer designed against a synthetic random sequence (non-existing in the mouse genome – green bar), inserted upstream the middle arm, allows genotyping of the animals. (**b**) The directed mutagenesis was confirmed by Sanger sequencing of the third exon in WT and homozygous stargazin KI^VL/VL^ animals. (**c**) Cresyl violet staining of brain slices from WT and stargazin KI^VL/VL^ animals showed no gross differences. Scale bar represents 1000 μm for lower magnification images and 200 μm for magnified images. (**d**) The brain expression pattern of stargazin in WT and stargazin KI^VL/VL^ animals was assessed by immunohistochemistry. Scale bar represents 1000 μm. (**e**) Total stargazin levels in the whole brain, hippocampal and cortical lysates from WT and stargazin V143L KI animals were evaluated by Western blot. See also Fig. S5.

To assess whether the V143L stargazin mutation affects stargazin protein levels and distribution across the brain, we performed immunolabeling of stargazin in brain coronal and sagittal slices from WT and stargazin V143L KI mice. Stargazin is broadly expressed throughout the mouse brain with high expression levels in the cerebral cortex, hippocampus, and cerebellum ^4^. Within the hippocampus, stargazin immunoreactivity was more intense in the *stratum oriens* of the CA1, CA2 and C3 regions, the *stratum lacunosum moleculare* of the CA1 and CA2 regions, and particularly in the *subiculum* (Fig. S5a,b). Stargazin immunoreactivity was similar in all genotypes (Fig. 3d and S5b,c), indicating that the expression of the ID-associated form of stargazin does not affect the protein brain-wide distribution and total expression levels. This was also confirmed by western blot analyses, using total lysates from the whole brain, cortex and hippocampus (Fig. 3e, Fig. S5d,e). The expression levels of stargazin and other TARPs were also assessed by qPCR and no changes were detected in samples from the cortex and hippocampus of KI mice, compared to WT littermates (Fig. S5f).

Since the V143L stargazin mutation was found in an ID patient ^16^, we asked whether stargazin V143L KI mice display alterations in motor function, anxiety-like behavior, cognitive and/or social performance that correlate with ID-like symptomatology. We began by assessing motor behavior in the open field test (Fig. S6a) and found that, whereas male stargazin KI^+/VL^ and KI^VL/VL^ mice showed comparable distance travelled and instant speed to WT male mice (Fig. S6c,e), female stargazin KI^VL/VL^ mice travelled longer distances, and stargazin KI^+/VL^ female mice showed higher instant speed than WT female mice (Fig. S6b,d), suggesting hyperactivity in female stargazin V143L KI animals. However, stargazin V143L KI mice did not display anxiety-like behaviors either in the open field (Fig. S6f) or in the elevated plus maze (Fig. S6g,h) tests, nor did they show depressive-like behavior in the forced swimming test (Fig. S6i,j). In an object displacement test for spatial memory evaluation, while WT animals preferred to spend time engaging with the displaced object, neither stargazin KI^+/VL^ nor KI^VL/VL^ mice showed this preference, and stargazin KI^VL/VL^ mice spent significantly less time exploring the object that was moved when compared to WT animals (Fig. 4a,b). Furthermore, male stargazin V143L KI animals failed to alternate above chance level in the T-maze spontaneous alternation test (Fig. S6k,l). In the contextual fear conditioning test for associative memory, stargazin KI^VL/VL^ mice presented less freezing behavior than WT animals (Fig. 4c,d). These observations suggest that the V143L mutation in stargazin elicits learning and memory impairments. Given the high expression of stargazin in the cerebellum (Tomita et al 2003), we assessed motor learning of stargazin V143L KI animals in the rotarod test (Fig. 4e-g). No significant motor abnormalities were displayed by mutant mice in the rotarod test (Fig. 4f), but whereas WT and stargazin KI^+/VL^ mice improved their performance the second day they were placed in the apparatus, stargazin KI^VL/VL^ mice failed to do so, suggesting an impairment in motor learning (Fig. 4g).

**Figure 4.**
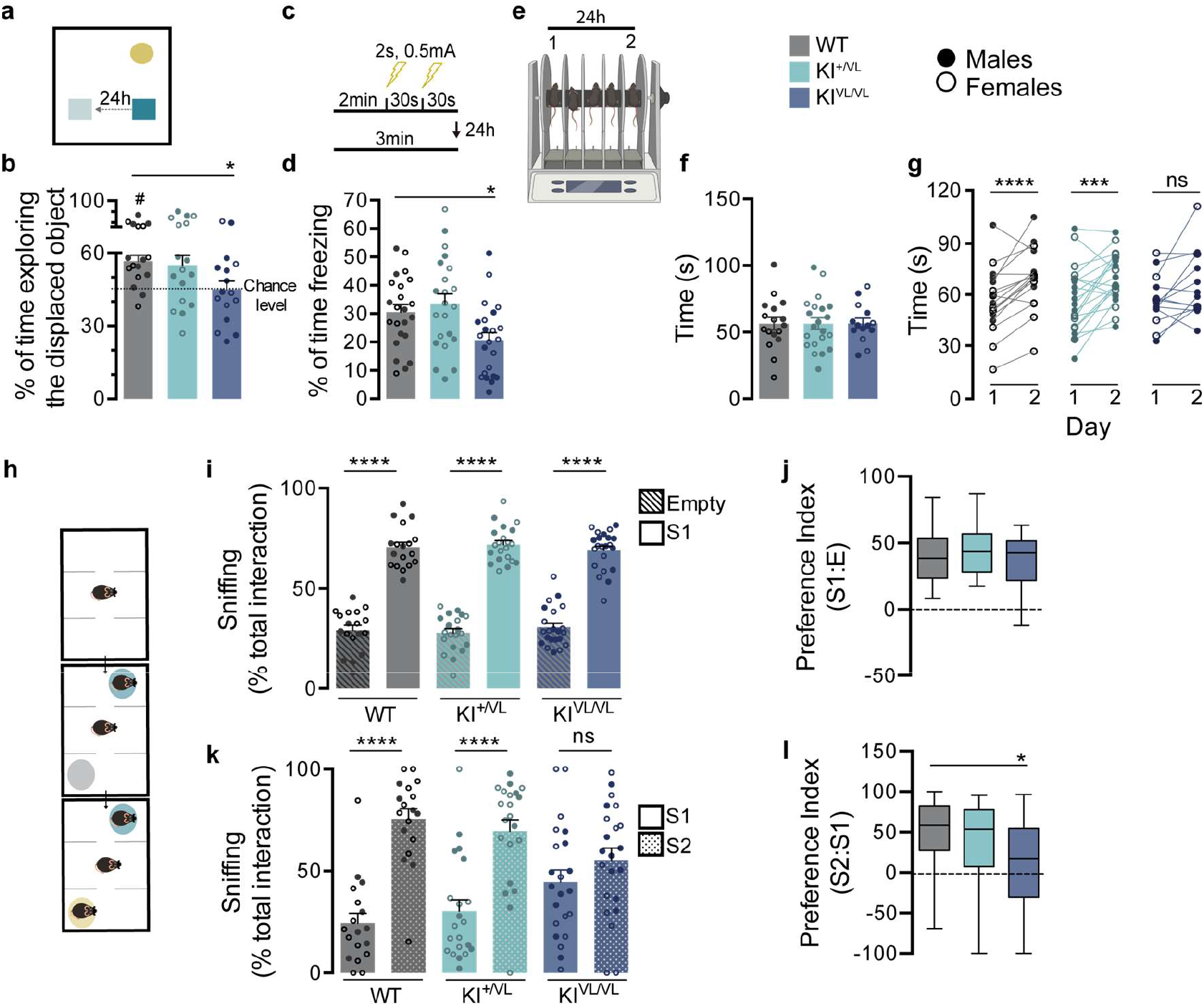
Stargazin V143L KI mice show cognitive and social deficits. (**a, b**) When subjected to the object displacement recognition test, homozygous stargazin V143L (KI^VL/VL^) mice spent less time exploring the displaced object when compared to their WT counterparts, and did not have preference for the displaced object. Data are presented as mean ± SEM. \**p* < 0.05, One-way ANOVA followed by Dunnet’s multiple comparison post hoc test, #*p*< 0.05, One-sample t-test to the value of 50%. n ≥ 16 (males and females) for all genotypes. (**c**,**d**) Homozygous stargazin V143L knock-in mice presented significantly less freezing behavior than WT counterparts in the contextual fear conditioning test. Data are presented as mean ± SEM. \**p* ≤ 0.05, One-way ANOVA followed by Dunnett’s multiple comparison post-hoc test. n ≥ 22 (males and females) for all genotypes. (**e**) Motor function and learning were evaluated using the rotarod test. (**f**) The average time spent on the rotarod did not significantly vary between genotypes. (**g**) Both WT and heterozygous stargazin V143L (KI^+/VL^) animals performed significantly better in the second day, whereas stargazin KI^VL/VL^ mice failed to show motor learning. \*\*\**p* < 0.001, \*\*\*\**p* < 0.0001, Ratio paired t-test, n ≥ 14 for all genotypes (males and females). (**h**) Mice were submitted to the three*-*chamber social interaction paradigm. The time spent approaching the cages, with and without the stranger stimulus mouse, was evaluated for 10 and 5 minutes, respectively. (**i**,**j**) All animals displayed social preference, but (**k**,**l**) stargazin KI^VL/VL^ mice showed no preference for a new stranger mouse in the arena, unlike WT and heterozygous stargazin V143L mice. Data are presented as mean ± SEM (**i, k**) and median with range (**j, i**). (**i, k**) \*\*\*\**p* < 0.0001, Two-way ANOVA followed by Sidak0s multiple comparison post-hoc test. n ≥ 13 (males and females) for all genotypes. (**j, i**) *p ≤ 0.05, Kruskal-Wallis followed by Dunn’s multiple comparison post-hoc test. n ≥ 13 (males and females) for all genotypes. See also Fig. S6.

Typically, ID patients display deficits in several social skills, including the will/ability to socially engage with other people. To determine whether stargazin V143L KI mice display social interaction deficits, we tested these animals in the three-chamber test. Stargazin V143L KI mice showed preference for a conspecific (Stranger 1) over an empty cage (E), similarly to WT mice (Fig. 4h-j). However, in the presence of a novel social partner (Stranger 2), contrarily to WT and stargazin KI^+/VL^ mice, stargazin KI^VL/VL^ mice did not prefer to interact with the unfamiliar animal (Fig. 4h,k,l). This result suggests a possible deficit in social recognition and/or alterations in the motivation for social novelty. The innate social behavior of nest building was not perturbed in stargazin V143L KI mice (Fig. S6n,o). Together, our results show that the ID-associated mutation in stargazin elicits cognitive and social deficits reminiscent of ID-like symptoms.

### Stargazin V143L mutant mice exhibit early hippocampal synaptic transmission defects

To assess whether the decrease in surface AMPA receptor levels observed *in vitro* in neurons expressing stargazin V143L has an impact in glutamatergic transmission *in vivo*, we performed whole-cell patch-clamp recordings in CA1 pyramidal neurons from acute hippocampal slices of P15-P20 stargazin V143L KI mice, to measure AMPA receptor-mediated miniature excitatory post-synaptic currents (mEPSCs). We found that the frequency of mEPSCs events was significantly decreased in neurons from stargazin KI^+/VL^ and KI^VL/VL^ mice compared to WT littermates (Figs. 5a,c). Interestingly, no changes in the amplitude of mEPSCs (Figs. 5a,b) or in the kinetics of these events (Fig. 5a) were observed. These data indicate that the V143L mutation in stargazin does not impact AMPAR gating, but affects synaptic transmission in the hippocampus.

**Figure 5.**
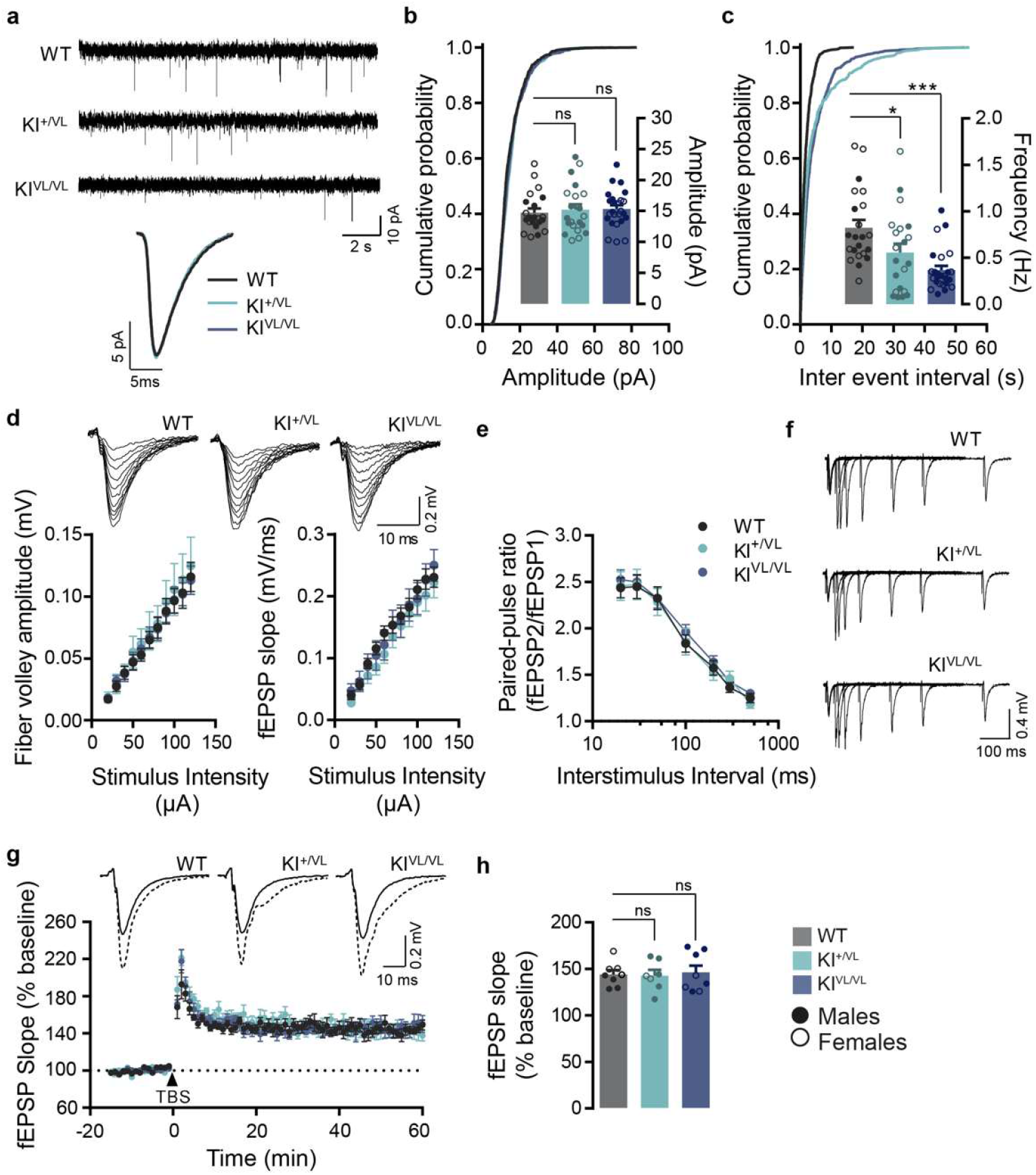
Decreased frequency of AMPAR-mediated mEPSCs in CA1 pyramidal neurons in stargazin V143L KI mice. **(a)** Representative traces of mEPSCs recordings and single average event of CA1 pyramidal neurons in acute hippocampal slices from WT, stargazin KI^+/VL^ and stargazin KI^VL/VL^ mice. Cumulative probability distribution and average mEPSCs amplitude (**b**) and frequency (**c**) plots, showing a reduction in frequency but not amplitude of mEPSCs in stargazin V143L KI mice (P15-P20). Data are presented as mean ± SEM. \**p* < 0.05, \*\*\**p* < 0.001, ns, not significant, Kruskal-Wallis followed by Dunn’s multiple comparison post-hoc test. n = 22 cells/10 animals (4 males and 6 females) for WT mice, n = 21 cells/8 animals (4 males and 4 females) for KI^+/VL^ mice, n = 25 cells/6 animals (3 males and 3 females) for KI^VL/VL^ mice. (**d**) Representative traces of evoked fEPSPs at CA1 synapses on hippocampal slices from P15-P20 WT and stargazin V143L KI mice, upon stimulation of CA3 pyramidal neurons Schaffer collaterals. Plots show the mean ± SEM of fiber volley amplitude and fEPSP slopes. Input-output curves were similar for all genotypes, indicating that evoked basal transmission in this synapse is not impaired in stargazin V143L KI mice. n = 17 slices/10 animals (5 male and 5 female) for WT mice, n = 11 slices/8 animals (4 male and 4 female) for stargazin KI^+/VL^ mice, n = 15 slices/10 animals (6 male and 4 female) for stargazin KI^VL/VL^ mice. (**e**) Paired-pulse facilitation at SC-CA1 synapses in stargazin V143L KI mice was not altered. Data are presented as mean ± SEM. n = 8 slices/6 animals (3 males and 3 females) for WT mice, n = 7 slices/5 animals (3 males and 2 females) for stargazin KI^+/VL^ mice, n = 8 slices/5 animals (3 males and 2 females) for stargazin KI^VL/VL^ mice. (**f**) Representative traces of paired-pulse stimulation evoked fEPSPs in WT and stargazin V143L KI mice. (**g**) LTP induced by theta burst stimulation (TBS) at SC-CA1 synapses was comparable between genotypes. Insets show representative traces of evoked fEPSPs before (solid lines) and after (dashed lines) LTP induction. Data are presented as mean ± SEM. n = 8 slices/6 animals (3 males and 3 females) for WT mice, n = 7 slices/5 animals (3 males and 2 females) for stargazin KI^+/VL^ mice, n = 8 slices/5 animals (3 males and 2 females) for stargazin KI^VL/VL^ mice. (**h**) Average fEPSP slope in the last 10 min of the recording post LTP-induction. Data are presented as mean ± SEM. ns, not significant, One-way ANOVA followed by Dunnett’s multiple comparison post-hoc test.

We next investigated the consequences of the ID-associated stargazin mutation in hippocampal functional connectivity and synaptic plasticity by recording field excitatory post-synaptic potentials (fEPSPs) in CA1 while stimulating the Schaffer collateral fibers. First, we tested the impact of the V143L stargazin mutation on the CA3-to-CA1 Schaffer collateral synaptic transmission and did not find significant differences in input/output curves (Fig. 5d). This suggests that the connectivity between CA3 pre-and CA1 post-synaptic sites is preserved in the hippocampus of stargazin V143L KI animals. Additionally, no significant alterations were found in the fiber volley amplitude between genotypes, indicating that there are no gross presynaptic impairments in stargazin V143L KI mice at these synapses (Fig. 5d). Indeed, when paired-pulse facilitation, a short-term strengthening of synaptic transmission, was assessed no overt alterations were observed (Fig. 5e,f), further supporting that the presynaptic function is intact in the hippocampus of stargazin V143L KI mice. Finally, we induced LTP in acute hippocampal slices using a theta burst stimulation protocol. Stargazin V143L KI mice showed normal LTP at Schaffer collateral-CA1 synapses (Fig. 5g,h). Altogether, these data indicate that, despite the decreased frequency of mEPSC events detected in CA1 pyramidal neurons (Fig. 5a,c), synaptic connectivity and theta burst-induced long-term synaptic potentiation in the Schaffer collateral-CA1 synapse are not severely impaired in stargazin V143L KI mice.

### Stargazin V143L mutant mice have reduced mature spine density on basal dendrites of CA1 hippocampal neurons

Dendrites and spines are key neuronal structures in signal integration and transmission. Changes in dendritic branch complexity and length, and spine density, volume and shape have been described in the brains of patients with neuropsychiatric disorders [reviewed in ^36^]. To understand if the stargazin V143L mutation impacts neuronal morphology, we performed Sholl analysis of CA1 pyramidal neurons of stargazin V143L KI mice. This structural analysis allows the evaluation of alterations in dendritic complexity based on the number of dendritic intersections at various distances from the soma. For that, we intravenously injected AAV9.hSyn.GFP in the mice to achieve sparse, Golgi-like, labelling of neurons and outline the morphology of dendrites and dendritic spines (Fig. S7a,b). Our results show that the stargazin V143L mutation has no impact on dendritic arbor complexity (Fig. S7c,d). Moreover, no statistically significant differences were found in the total dendritic length of basal and apical dendrites (Fig. S7e).

Next, we evaluated the effects of the stargazin V143L mutation on dendritic spine density and morphology in the hippocampus of stargazin V143L KI mice. While no significant differences in total dendritic spine density were observed, there was a significant decrease in the density of mature spines, namely mushroom and stubby spines, on basal dendrites of CA1 pyramidal neurons from stargazin KI^+/VL^ and KI^VL/VL^ mice when compared to WT littermates (Fig. 6a,b). Additionally, stargazin KI^+/VL^ and KI^VL/VL^ mice displayed increased density of branched spines and of immature spines (thin and filopodia) on basal dendrites (Fig. 6a,b). In contrast, no changes in spine density or morphology were found on apical dendrites of CA1 pyramidal neurons of stargazin V143L KI mice (Fig. 6c,d). Overall, there was a significant decrease in the percentage of mature spines on basal dendrites of stargazin V143L KI mice neurons, whereas no changes were observed on the morphology of spines on apical dendrites (Fig. 6e). Interestingly, in the CA1 region stargazin was more expressed in the *stratum oriens*, where the basal dendrites are located, when compared to the *stratum radiatum*, where apical dendrites are placed (Fig. S5a). This differential pattern of stargazin expression within the CA1 region correlates with the pronounced effects in spine maturation observed on basal dendrites. To conclude, these data show that the stargazin V143L mutation specifically results in spine dysmorphogenesis on basal dendrites without having a major role on the formation/elimination of dendritic spines.

**Figure 6.**
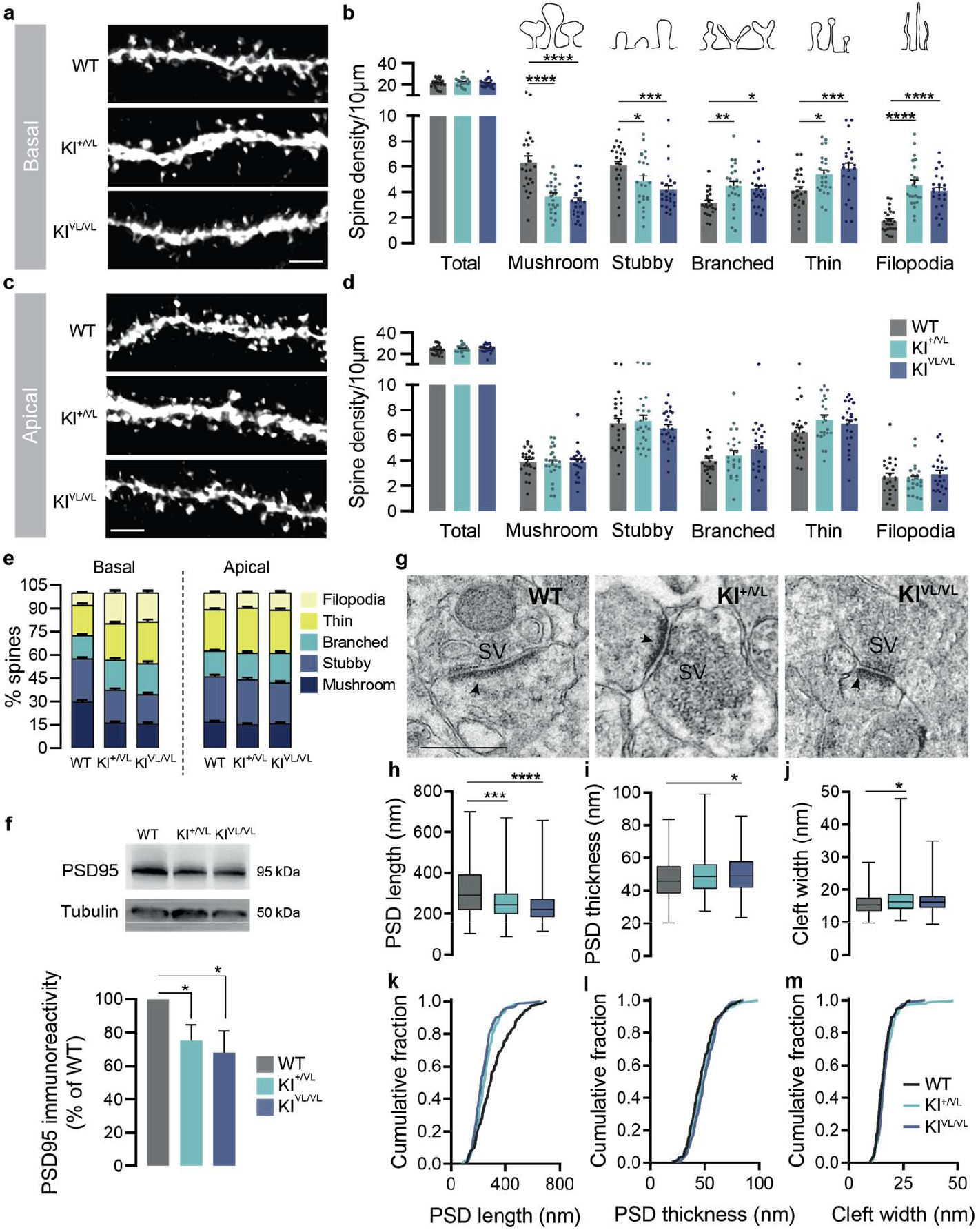
Stargazin V143L KI mice present alterations in hippocampal spine morphology and PSD ultrastructure. (**a-e**) Although no changes were observed in the density of spines from both basal and apical dendrites in CA1 neurons, stargazin V143L KI animals presented a decrease in mushroom and stubby spines and an increase in branched, thin and filopodia spines, in basal dendrites, indicating a defect in spine maturation. Data are presented as mean ± SEM (**b, d**) or as percentage of the total number of spines (**e**). \**p* < 0.05, \*\**p* < 0.01, \*\*\**p* < 0.001, \*\*\*\**p* < 0.0001, Two-way repeated measures ANOVA followed by Dunnet’s multiple comparison post-hoc test. n = 24 branches/3 animals for all genotypes. Scale bar represents 2 μm. (**f**) Immunostaining of PSD95 by Western blot showed a significant decrease of its levels in hippocampal samples from stargazin V143L KI mice compared to WT controls. \**p* < 0.05, One-sample t-test to the value of 100%. The staining of tubulin of hippocampi samples is the same presented in Figure 3e. (**g**) Representative electron transmission microscopy images of hippocampal synapses from WT, stargazin KI^+/VL^ and stargazin KI^VL/VL^ animals, and (**h**,**k**) quantification of post-synaptic density length, (**i**,**l**) thickness and (**j**,**m**) cleft width. Data are presented as median with range. \**p* < 0.05, \*\*\**p* < 0.001, \*\*\*\**p* < 0.0001, Kruskal-Wallis followed by Dunn’s multiple comparison post hoc-test. n = 153 PSDs/2 animals for WT mice, n = 203 PSDs/2 animals for KI^+/VL^ mice, n = 236 cells/2 animals for KI^VL/VL^ mice. The arrows indicate post-synaptic densities. SV, synaptic vesicles. Scale bar represents 500 nm. See also Figs. S7 and S8.

To further explore the alterations in spine morphology, we performed ultrastructural analysis of the post-synaptic density (PSD) in hippocampal spines from WT and stargazin V143L KI mice using electron microscopy. Our analysis uncovered a significant decrease in the length of PSDs from stargazin KI^+/VL^ and KI^VL/VL^ mice (Fig. 6g,h,k), as well as an increase in the thickness of PSDs from stargazin KI^VL/VL^ mice when compared with WT littermates (Fig. 6g,i,l), highlighting potential alterations in post-synaptic structure and composition. In stargazin KI^VL/VL^ mice there was a slight but significant increase in the synaptic cleft width (Fig. 6j,m). Moreover, the total levels of PSD95 in the hippocampus of stargazin V143L KI mice were significantly reduced compared to WT littermates (Fig. 6f). PSD95 has an important role in silent synapse maturation ^37^ and PSDs with smaller size and decreased PSD95 content are less stable ^38^. The decreased PSD95 levels further support an impairment in spine maturation in the hippocampus of stargazin V143L KI mice. Since stargazin is also expressed in the cortex, we analyzed the effects of the stargazin V143L mutation in the ultrastructure of PSDs from this brain region as well (Fig. S8a-g). We also observed a significant decrease in the length of PSDs of stargazin KI^VL/VL^ mice (Fig. S8a,b,e). However, in contrast to what was observed in the hippocampus, there was a significant decrease in the thickness of PSDs from stargazin KI^+/VL^ mice (Fig. S8a,c,f). No changes in the synaptic cleft width were observed in cortical PSDs (Fig. S8a,d,g) nor in the total levels of PSD95 (Fig. S8h) in the cortex of stargazin V143L KI mice.

Together, these data reveal that the stargazin V143L mutation leads to an increase in the density of spines with immature morphology and to ultrastructural changes in post-synaptic compartments, indicating a general spine immaturity state in certain hippocampal subregions of stargazin V143L KI mice. Combined with our functional characterization showing decreased frequency of mEPSC events in CA1 neurons from stargazin mutant mice, this strongly suggests that the stargazin V143L mutation perturbs spine maturation and diminishes functional synaptic contacts in specific hippocampal subcircuits.

### Stargazin phosphorylation and interaction with GluA1 are decreased in stargazin V143L mutant mice

In a final set of experiments, we sought to determine whether the dendritic spines immaturity (Fig. 6a,b,e) and the decreased PSD length (Figs. 6h-m and S8a,b,e) found in the brain of stargazin V143L KI animals are accompanied by altered composition of the PSDs. We isolated PSDs from the cerebral cortex of WT, stargazin KI^+/VL^ and KI^VL/VL^ littermate mice (Fig. S8i) and quantified their content in stargazin, GluA1 and GluA2 AMPAR subunits, as well as PSD95 (Fig. 7a-e). Stargazin expression was decreased in the PSDs of stargazin KI^+/VL^ and KI^VL/VL^ mice compared to WT mice (Fig. 7a, despite not significantly changed total expression levels of stargazin in mutant mice -Fig. 3e), in agreement with increased cell surface mobility and decreased synaptic residence of stargazin V143L compared to the WT protein (Fig. 2a-f). Heterozygous stargazin V143L mice showed decreased levels of GluA1, GluA2 and PSD95 at the PSD, which were not significantly changed in homozygous stargazin KI^VL/VL^ mice (Fig. 7a, c-e). These observations suggest that the V143L mutation in stargazin impairs its expression at the synapse, which interferes with the synaptic content of AMPAR subunits. However, in homozygous mutant mice other TARPs may preferentially bind to AMPAR subunits and partially compensate for mutant stargazin in AMPAR synaptic trafficking. Indeed, and in agreement with our molecular dynamics analyses (Fig. 1e), immunoprecipitation of stargazin V143L from the cerebral cortex of stargazin KI^VL/VL^ mice showed decreased co-immunoprecipitation of GluA1, compared with stargazin immunoprecipitated from the cortex of WT littermate mice (Fig. 7f,g), indicating that the V143L mutation in stargazin impairs its interaction with AMPAR subunits *in vivo*.

**Figure 7.**
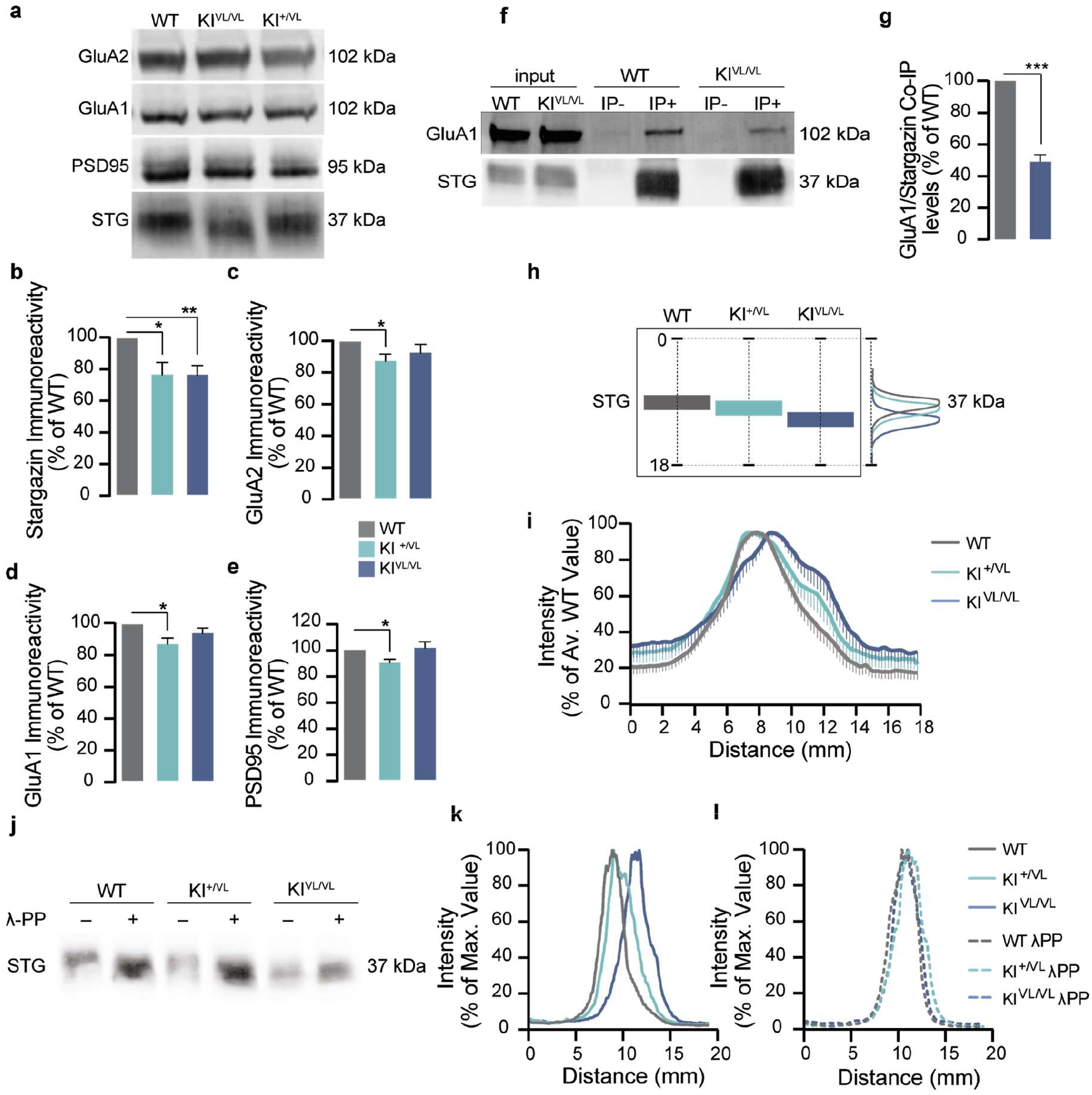
Stargazin V143L expression modifies the composition of PSDs. (**a**) Western blot analysis of cortical PSDs (see also Fig. S8j) showed that heterozygous stargazin KI^+/VL^ animals and homozygous stargazin KI^VL/VL^ mice have decreased synaptic levels of stargazin (**b**). GluA2 (**c**), GluA1 (**d**) and PSD95 (**e**) levels were also reduced in the PSDs of stargazin KI^+/VL^ animals. Data were normalized for WT values and are presented as mean ± SEM. One-sample t-test to the value of 100%. * *p* ≤ 0.05, \*\**p* < 0.01, n ≥ 6 for all conditions. (**f**) Immunoprecipitation of stargazin from the cortices of WT and stargazin KI^VL/VL^ animals. (**g**) Co-immunoprecipitation of GluA1 with stargazin was significantly reduced in the KI^VL/VL^ cortices. Data were normalized for immunoprecipitated stargazin in each respective condition. Data normalized to WT values and presented as mean ± SEM. One-sample t-test to the value of 100%, \*\*\**p* < 0.001, n = 5 for all conditions. (**h**) The SDS-PAGE migration pattern of stargazin from WT, stargazin KI^+/VL^ and KI^VL/VL^ mouse cortical PSD extracts (**a**) was analyzed by quantifying the distribution of the intensity of the bands along the length of the lane. (**i**) In PSDs isolated from stargazin KI^VL/VL^ animals, stargazin migrated faster in the SDS-PAGE. Data were normalized for the average maximum intensity of WT stargazin for each western blot membrane and are presented as mean ± SEM, n = 9. (**j**) Representative images from stargazin labelling profile in cortical PSD samples isolated from WT, stargazin KI^+/VL^ and KI^VL/VL^ samples non-treated (**k**) or treated (**l**) with λ-Phosphatase. λ-Phosphatase treatment of isolated cortical PSDs induced a shift in the apparent molecular weight of stargazin in WT and stargazin KI^+/VL^ PSD samples, but not in stargazin KI^VL/VL^ PSD samples, suggesting deficient phosphorylation of the stargazin V143L protein variant detectable in homozygous stargazin V143L KI animals.

The function of stargazin is regulated by the phosphorylation of serine residues in the cytoplasmic C-terminal tail of the protein ^12^, and these phosphorylation events regulate stragazin interaction with membrane lipids ^39^, its binding to PSD95 ^9,40^, and the diffusional trapping of AMPARs at synaptic sites ^33^. The migration pattern of stargazin in denaturing SDS-PAGE conditions correlates with the phosphorylation state of the protein, with phosphorylated stargazin showing slower migration in SDS-PAGE ^11,12^. We noticed that, in PSDs isolated from the cortex of stargazin KI^VL/VL^ mice, stargazin showed faster migration compared to PSDs isolated from WT mice, whereas an intermediate migration pattern was detected in PSD isolated from heterozygous stargazin V143L mice (Fig. 7h,i). To test for altered phosphorylation of mutant stargazin, we treated cortical extracts with λ-phosphatase before PSD purification and found that stargazin bands in PSDs isolated from the cortex of WT or heterozygous stargazin KI^+/VL^ mice shifted to a lower apparent molecular weight, putatively corresponding to the unphosphorylated form of the protein ^11,12^ and coincident with the stargazin band in PSDs isolated from untreated cortical extracts from stargazin KI^VL/VL^ mice (Figs. 7j-l). In fact, the gel mobility of the stargazin band in PSDs isolated from stargazin KI^VL/VL^ mice was unchanged by λ-phosphatase treatment (Figs. 7j-l), indicating that the protein is in a dephosphorylated form. These findings are consistent with decreased phosphorylation of V143L stargazin.

## DISCUSSION

In this study, we employed molecular dynamics analyses, *in vitro* and *in vivo* models to study how an ID-associated mutation in the third transmembrane domain of stargazin impacts the AMPAR:stargazin complex, hippocampal synaptic architecture, synapse function and behavior. Our data suggest that the V143L mutation in stargazin critically affects stargazin interaction with the AMPAR complex, leads to decreased stargazin phosphorylation, decreases AMPAR-mediated synaptic transmission and contributes to spine immaturity in CA1 hippocampal neurons. A striking aspect of our study is that it reveals not only the pathogenic effect of a mutant form of stargazin associated with disease, but also unveils critical roles for stargazin in regulating synapse structure and function in the hippocampus and in shaping cognitive and social behavior.

Structural analyses highlight that the main interface between AMPAR:stargazin is in the TMD of the complex, where the ECD of stargazin acts as a scaffold on the receptor. This interaction is mainly mediated by TMD3 and TMD4 from stargazin, M1 and M2 from *Main GluA*, and M4 from *Auxiliary GluA*. Our analysis showed a positively correlated motion between these substructures (Fig. 1e), which presented lower ΔSASA values, indicating their higher contribution to the overall interface. The stargazin V143L mutation hinders the correlated motion between TMD regions of stargazin and GluA, especially at M4 of *Auxiliary GluA*. Furthermore, ΔSASA values for these substructures are higher in the systems containing mutated stargazin. X site shows greater differences between WT and mutant stargazin, suggesting that the stargazin V143L mutation especially weakens AMPAR:stargazin interaction at the X site. In agreement with a weakening effect of the V143L mutation on the interaction between stargazin and AMPAR, we found a decreased co-immunoprecipitation of GluA1 with stargazin V143L (in cortical brain lysates from stargazin KI^VL/VL^ mice) compared to WT stargazin (Fig. 7f,g). Furthermore, we observed decreased trafficking of AMPAR to the synapse upon deletion of stargazin, which was not rescued upon expression of the ID-associated variant of stargazin in cortical neurons (Fig. 2g-l).

To date, the physiological roles of TARPs have been studied using knock-out mice for the different TARPs, alone or in combination [reviewed in ^2^]. These analyses have provided crucial insight into partially overlapping although non-redundant functions for different TARPs, but are hindered by possible compensatory effects that may arise in the absence of the endogenous proteins. Examining knock-in mouse models expressing mutant forms of stargazin associated with disease has the double advantage of informing on the endogenous role of stargazin, by analyzing the effects of loss of function mutant variants which are still expressed, and on possible pathogenic mechanisms elicited by human stargazin mutations. In this study, we have found that the V143L mutation in stargazin triggers a striking decrease in the frequency of mEPSCs in hippocampal CA1 pyramidal neurons (Fig. 5a-c), and leads to a decrease in spine maturity in CA1 basal dendrites (Fig. 6a, b,e), and to ultrastructural alterations in the post-synaptic compartment, in particular a significant reduction of the PSD length (Fig. 6g-m). These observations suggest that despite the expression of multiple other TARP members in the hippocampus, including γ3 and high enrichment in γ8 (Tomita et al 2003), stargazin is required for normal spine and PSD development and for maintaining a full complement of functional synapses. Our results are in line with experiments using *stargazer*/*γ8*-knock-out mice, which showed that AMPAR-mediated transmission in CA1 pyramidal neurons is further reduced, compared to the reduction observed in *γ8*-knock-out mice ^41^, despite the fact that CA1 pyramidal neurons from *stargazer* mice did not show alterations in the ratio of AMPA to NMDA EPSC amplitudes ^42^. The synergistic reduction in AMPAR-mediated transmission in the *stargazer/γ8* double knock-out mice implies some degree of functional redundancy for the two TARPs. If mutated stargazin is expressed, its incorporation in AMPAR complexes, even if less efficient than WT stargazin, will thus exert pathogenic effects, as suggested by the reduction in the frequency of mEPSCs and in spine maturation that we observed in stargazin V143L mice. These results are also in agreement with electron microscopy data showing that at Schaffer collateral/commissural synapses in the CA1 hippocampal region the presence of stargazin correlates with higher density of AMPAR expression ^43^ and thus presumably with the presence of a higher number of functional synapses.

Our data show a specific effect of the stargazin V143L variant in spine maturation in basal dendrites in CA1 neurons, which was not observed in apical dendrites (Figs. 6a-e). Indeed, despite the significant change in the frequency of mEPSC in CA1 neurons in stargazin V143L mice (Figs. 5a-c), we found no changes in field EPSC slope (Fig. 5d) or in LTP (Fig. 5g,h) in Schaffer collateral-CA1 synapses recorded in the *stratum radiatum*, which are located in the apical dendrites of CA1 pyramidal neurons. These observations indicate that stargazin has a specific role in maintaining spine structure in CA1 basal dendrites, which is in agreement with the higher expression levels of stargazin in the hippocampal *stratum oriens*, compared with the *stratum radiatum* (Fig. S5a). Altogether, our data suggest that, besides the well-described brain region-and cell type-specific roles of TARPs, there may be subcellular-specific roles that are determined by the subcellular distribution pattern of different TARPs.

The V143L variant of stargazin was found to be dephosphoryated in cortical PSDs isolated from homozygous stargazin V143L KI mice, compared to WT PSDs (Fig. 7h-l). Phosphorylation of stargazin in its C-terminal region disrupts electrostatic interaction between the membrane and stargazin C-tail ^39^, promotes the extension of the C-tail into the cytoplasm and binding to PSD95 ^40^, and triggers diffusional trapping of AMPARs at synaptic sites ^33^. Stargazin phosphorylation has been proposed to regulate Hebbian forms of synaptic plasticity ^12^ and to mediate experience-dependent plasticity and synaptic scaling ^10,11^. The lower level of phosphorylation of stargazin-V143L compared to the WT protein likely underlies its higher membrane diffusion rate at the membrane and its impaired capacity in supporting AMPAR synaptic traffic (Fig. 2). The low phosphorylation of stargazin V143L may also determine the sequestration of its C-terminal tail in the plasma membrane and thus impair it from undergoing liquid-liquid phase separation with PSD scaffold proteins ^9^. Changes in hippocampal spine maturation and in the ultrastructure of the hippocampal PSDs (Fig. 6) may thus be a consequence of defective stargazin V143L phosphorylation, and may be reflected in the decreased number of functional synapses detected in our mEPSC analyses in CA1 hippocampal neurons (Fig. 5a-c). While it is likely that the aberrant stargazin V143L phosphorylation contributes to the physiological effects observed, our MD analysis, which does not take into account post-translational modifications in stargazin, also suggests compromised function for the V143L stargazin variant.

In this study we found that the V143L mutation in stargazin in male and female stargazin KI^VL/VL^ mice leads to altered spatial memory (Fig. 4a,b), assessed using the object displacement test, as well as perturbed associative memory in the contextual fear conditioning test (Fig. 4c,d). These alterations in hippocampal-dependent cognitive behavior are likely to be related to the changes in mEPSC frequency, in spine maturity and in PSD ultrastructure that we identified in the hippocampus of these mice. We did not detect changes in social interaction in the three chamber test in stargazin V143L mice (Fig. 5i,j), but stargazin KI^VL/VL^ mice showed impairment in preference for social novelty (Fig. 5k,l), suggestive of either a perturbation in social memory or a lack of motivation for social novelty. Stargazin KI^VL/VL^ mice also displayed impaired motor learning in the rotarod (Fig. 5g), pointing to possible functional and structural alterations in the cerebellum caused by the stargazin V143L mutation. Given the elevated expression of stargazin in the cerebellum ^4^ and its non-redundant functions in cerebellar excitatory synapses in several cerebellum circuits ^2,7,42^, future studies should examine cerebellum circuit-specific dysfunction triggered by the ID-associated stargazin mutation. The cognitive and social behavioral dysfunctions displayed by stargazin V143L mice most likely arise from alterations in a combination of brain circuits, depending on the stargazin expression pattern and its synaptic roles in different cell types. Together, our data provide the first evidence for the causal implication of stargazin in the pathogenesis of neurodevelopmental disorders.

## METHODS

### Modeling the three-dimensional protein structure

The three-dimensional (3D) structure of stargazin was constructed by homology modelling using the MODELLER package ^44^, the target sequence retrieved from UniProt [^45^, Q9Y698] and two templates: GluA2:stargazin complex [PDB-ID: 6DLZ ^28^; electron microscopy with 3.9 Å resolution; Human Organism; 99.5% sequence similarity]. The best one hundred models from MODELLER ^44^ were evaluated by DOPE score, z-score ^46,47^, LGscore and MaxSub ^48^. The final model loops were further optimized. Due to the lack of a well-defined secondary structure with subsequent high conformational heterogeneity, the C-terminal of this protein, located at the intracellular level, was removed from the final model. The V143L stargazin mutation was built using the mutagenesis tool of PyMOL, creating the ID model. The 3D structure of GluA2 (Ligand-binding domain – LBD -and transmembrane domain – TMD) was also constructed using the MODELLER package ^44^ and the subunit of AMPAR as the template (PDB-ID: 6DLZ), with the sequence P42262 from UniProt. The final model was selected using the previous criteria and the complexes AMPAR:stargazin (WT and V143L variant) were obtained by the superimposition of the stargazin and AMPAR models with 6DLZ structure. The final model 3D structure is illustrated in Figure 1b.

### Molecular dynamics simulations

Molecular Dynamics (MD) simulations of AMPAR:stargazin WT and mutated form (V143L variant) were performed using GROMACS 2018.4 ^49^ and the CHARMM36 force field ^50^. The complex orientation in the membrane was obtained through the oriented crystal of GluA2:stargazin complex (PDB-ID: 6DLZ). Systems were built using CHARMM-GUI ^51,52^ membrane builder with a bilayer membrane of POPC:Cholesterol (9:1 ratio) to replicate the physiological environment. Each complex was solvated by a TIP3 water box and 0.15 M of NaCl. The final WT and mutated systems were constituted by 518.000 and 521.000 atoms, respectively.

The systems were subjected to an initial minimization to remove bad contacts using the steepest descent algorithm. Subsequently, they were heated using the Berendsen-thermostat at 310 K in the NVT ensemble over 7 ns, followed by an NPT ensemble of 20 ns with a semi-isotropic pressure coupling algorithm ^53^, which is used to keep the pressure constant of one bar. Long-range electrostatic interactions were treated by the fast smooth Particle-Mesh Ewald method ^54^. All bonds, involving hydrogen atoms within protein and lipid molecules were constrained using the linear constraint solver (LINCS) algorithm ^55^. Additionally, a cut-off distance of 12 Å was attributed to Coulombic and van der Waals interactions. Three independent replicas were run for each system during 0.5 μs, of which the first 0.15 μs of equilibration were left out of the further analysis.

Root mean square deviations (RMSD) calculations were performed using the Cα atoms by GROMACS package ^44^. The cross-correlation analysis (CCA), which tracks the movements of two or more sets of time series data relative to one another, was calculated by Bio3D R package ^56^ for residue-level dynamic analysis using the Cα trajectory. CCA analysis provides atomistic detail about the dynamic nature of proteins, and in particular allows the differentiation between regions that exhibit correlated or anticorrelated motions with others, in the same or in the opposite direction, respectively ^57^. The solvent-accessible surface area (SASA) analysis for each residue was performed using GROMACS package ^49^. These analyses were performed for the bound and unbound systems, and ΔSASA by residue was calculated as SASAAMPAR:STG – (SASAGluA2 + SASASTG). ΔSASA values, summed by substructure, provide another quantitative measure of conformational change upon protein coupling ^58^.

### Primary cortical neurons

Primary cultures of rat cortical neurons were prepared from the cortices of E17 Wistar rat embryos. Briefly after dissociation, the cortices were incubated with trypsin (0.06%, 10 min, 37°C, GIBCO Invitrogen) in Ca^2+^-and Mg^2+^-free HBSS (5.36 mM KCl, 0.44 mM KH2PO4, 137 mM NaCl, 4.16 mM NaHCO3, 0.34 mM Na2HPO4.2H2O, 5 mM glucose, 1 mM sodium pyruvate, 10 mM HEPES and 0.001% phenol red), washed 6 times with HBSS and then mechanically dissociated. After counted, the cells were plated, at a low density (0.3×10^6^ cells per 60 mm culture dish), in neuronal plating medium (MEM supplemented with 10% horse serum, 0.6% glucose and 1 mM pyruvic acid) in five poly-D-lysine (0.1 mg/ml) coated coverslips (18 mm). The medium was replaced, after 2 hours, by Neurobasal medium supplemented with SM1 (StemCell Technologies), 0.5 mM glutamine and 0.12 mg/ml gentamicin. Neurons grew facing a confluent feeder layer of astroglial cells but were kept apart from the glial cells by wax dots placed on the coverslips ^59^. The cultures were treated with 5 μM cytosine arabinoside, two days after plating, to prevent the overgrowth of glial cells and were maintained in an incubator with 5% CO2, at 37°C. Conditioned medium was partially replaced by fresh, SM1 supplemented neurobasal medium every 3 days. Primary cortical cultures were used for imaging.

### Transfection of cortical neurons

Neurons were transfected using a calcium phosphate-mediated transfection protocol ^60^. A CaCl2 solution (2.5 M in 10 mM HEPES) was added, dropwise, to the diluted DNA. This solution was then added to the equivalent volume of HEPES-buffered transfection solution (274 mM NaCl, 10 mM KCl, 1.4 mM Na2HPO4, 11mM dextrose and 42 mM HEPES, pH 7.2). The DNA precipitates were added, dropwise, to the coverslips in conditioned medium and 2 mM of kynurenic acid. The cultures were incubated for 2 h at 37°C and 5% CO2. The DNA precipitates were dissociated by incubating the cells with acidified medium, for 15 minutes at 37°C and 5% CO2. Coverslips were then transferred to the original astroglial-containing dish.

### Immunocytochemistry and imaging

In order to stain surface proteins, live cells were incubated with anti-GluA (MAB2263; Millipore) primary antibody diluted in conditioned medium for 10 minutes and fixed for 15 min in 4% sucrose/ 4% paraformaldehyde in PBS at room temperature. Following 3 washes with PBS, the cells were incubated with the secondary antibody (Molecular Probes) diluted in 3% BSA, in PBS, for 45 min, 37°C. After 6 washes with PBS, cells were permeabilized for 5 min with 0,25% Triton X-100, in PBS at 4°C. Unspecific staining was blocked by incubation with 10% (w/v) BSA in PBS for 30 min, at 37°C. In order to label PSD95 (MA1-045; Thermo Scientific) and MAP2 (ab5392; Abcam), neurons were incubated with the primary antibodies diluted in 3% BSA in PBS for 2h at 37°C or overnight at 4°C. Before and after incubating with the secondary antibodies, also diluted in 3% BSA in PBS, for 45 min, 37°C, cells were washed 6 times with PBS. Coverslips were mounted in DAKO fluorescent mounting medium. The imaging was performed using a Zeiss Axiovert 200 M microscope and a 63X (NA1.4) oil objective. Blind-to-condition quantification was performed in ImageJ analysis software, with a macro that automatized quantification steps. The region of interest (ROI) was chosen randomly, by using MAP2 and/or GFP staining to confirm that the selected dendrite was from a transfected neuron. The threshold was defined to include detectable clusters and the signal intensity of the particles of the selected area was analyzed. Synaptic puncta were defined by their colocalization with PSD95.

### Quantum dots labeling, imaging and analysis

Low-density 12 days *in vitro* (DIV) cells were co-transfected with plasmids encoding Homer-GFP, for synapse identification, and HA-tagged WT stargazin or the V143L stargazin variant. At DIV 14, cells were incubated for 10 min at 37°C with anti-HA antibody (3F10; Roche) (1:3000) diluted in conditioned medium. After one washing step, anti-rat IgG conjugated QD655 (diluted 1:10 in PBS) was diluted in conditioned medium with BSA 2% (1/2000) and added to cells for 5 min at 37°C. All washes were performed in ECS containing NaCl 145mM, KCl 5mM, Glucose 10mM, Hepes 10mM, CaCl2 2 mM and MgCl2 2mM, supplemented with BSA 2% at 37°C. Neurons were mounted in an open chamber (K.F. Technology SRL) and imaged in ECS. Single-particle tracking was performed as in (Opazo et al 2010). Cells were imaged at 37°C on an inverted microscope (Axio Observer Z1, Carl Zeiss) equipped with a Plan Apochromat 63X-1.4 numerical aperture oil objective. Homer1C-GFP signal was detected by using an HXP fluorescence lamp (For QDs: excitation filter 425/50 and emission filters 655/30, Chroma). Fluorescent images from QDs were obtained with an integration time of 50 ms with up to 600 consecutive frames. Signals were recorded with a digital CMOS camera (ORCA Flash 4.0, Hamamatsu). The tracking of single QDs was performed using the Metamorph and Matlab (Mathworks Inc., Natick, USA) software tools. Due to random blinking, the trajectories were not continuously tracked, instead, when the positions before and after the dark period were compatible with borders set for maximal position changes between consecutive frames and blinking rates, the subtrajectories of the same molecule were reconnected. MSD curves were calculated for reconnected trajectories of at least 20 frames. The QDs were considered synaptic if colocalized with Homer-1c dendritic clusters for at least five frames. Diffusion coefficients were calculated by a linear fit of the first 4–8 points of the mean square displacement (MSD) plots versus time depending on the length of the trajectory within a certain compartment. The resolution limit for diffusion was 0.0075 μm^2^/s as determined by ^61^, whereas the resolution precision was ∼40 nm.

### Animal generation and maintenance

Stargazin V143L KI mice were generated by inserting a single nucleotide mutation in the third exon of the *Cacng*2 gene. The targeting vector was introduced through homologous recombination in R1 cells, as described previously ^62^. Mice were viable and born at the expected Mendelian ratio. Genotyping was performed by PCR from mouse ear or tail DNA using a forward primer for the WT allele (AAGGGACCCTCCGTCCTCTC), a forward primer for the KI allele (GGGCCCGGTGCAATACACGC) and a reverse primer for both the reactions (CATCGGGCATGGATCCTCAGTTC). Mice were maintained at 22°C and 60% humidity under a 12h light/dark cycle. Food and water *ad libitum* were provided. During this study, both male and female animals were used. The imaging, biochemical and behavioral analyses were performed in mice with 8 to 10 weeks and electrophysiology recordings were performed in 15-20 days-old animals. All the procedures involving animals were performed according to the guidelines established by the European Union Directive 2010/63/EU and the experiments were previously approved by the institutional animal welfare body (ORBEA) and the national competent authority (DGAV).

### Nissl Staining

Eight-week old mice were anesthetized with isoflurane and perfused with ice cold PBS followed by 4% paraformaldehyde in PBS. Whole brains were kept in 4% paraformaldehyde in PBS overnight and then transferred to a 30% sucrose in PBS solution for at least 24 hours. Brains were sliced in the cryostat (Thermo Cryostar NX50, Thermo Fisher Scientific, USA) to obtain 50 μm coronal slices which were mounted in gelatin-coated slides. The slides were briefly washed with water and then submersed in a cresyl violet solution, for 5 minutes. After 2 washes with water, the slices were decolored with 100% ethanol for 2 minutes and incubated for 2 minutes in xylene before mounting with Permount mounting medium (Fisher scientific). Brain slices were digitized using a Zeiss Axio Scan.Z1 slide scanner (Carl Zeiss, Germany) equipped with a Plan Apochromat 20X-0.8 numerical aperture air objective.

### Immunohistochemistry

For stargazin immunofluorescence staining, 50 μm coronal and sagittal brain slices were prepared as described above. Free-floating sections were rinsed 3 times in PBS for 10 minutes and then permeabilized and blocked for 1 hour at room temperature with 0.25% Triton X-100 and 5% goat serum in PBS. After that, slices were incubated overnight at room temperature with the primary antibody (AB_2571844, Frontier Institute Co., Japan) diluted 1:200 in 0.25% Triton X-100 and 2% goat serum in PBS, followed by 3 washes with 0.25% Triton X-100 in PBS. Sections were incubated with the secondary antibody (anti-rabbit Alexa Fluor 568, Molecular Probes, USA) diluted 1:500 in 0.25% Triton X-100 and 2% goat serum in PBS, at room temperature for 2 hours. Nuclei were visualized by staining with 1μg/mL Hoechst 33342 in PBS for 5 minutes at room temperature. Lastly, after 3 washes of 10 minutes in PBS, the sections were mounted in gelatinized slides using Dako mounting medium (Glostrup, Denmark). Images were acquired on a Carl Zeiss Axio Imager Z2 upright widefield microscope (Carl Zeiss, Germany) using a Plan-Apochromat 20x air objective (NA 0.8) or in an LSM 710 Confocal microscope (Zeiss, Germany) with a Plan Apochromat 63x (NA 1.4) oil objective.

## Behavior analyses

### Object displacement test

The ODT was performed in a 40×40 cm open field arena. The test consisted of five trials. In the first trial the animals acclimatized to the empty arena for 6 minutes. In the three following trials, the animals were allowed to explore, for 6 minutes, two different objects located in a fixed position. In the fifth trial, conducted 24 hours later, one of the objects was displaced and the time spent exploring the non-displaced and the displaced object was evaluated.

### Contextual fear conditioning test

The contextual fear conditioning test was performed in an electrified wire-bottom 20×20 cm cage. On the first day, the animal was placed for 2 minutes in the cage before receiving a 2s-long foot shock of 0.5 mA. After 30 seconds, a second foot shock with the same magnitude and duration was administered, another 30 seconds later the animal was removed from the cage. After 24h, the animal was placed in the same cage and it was recorded for 3 minutes. The time spent in freezing behavior was scored using the Observer XT 12 software (Noldus, Netherlands).

### Rotarod test

Motor function and learning was evaluated using the accelerated rotarod (Med Associates), 4 to 40 rpm in 5 min. The time withstood in the rotating beam in three successive trials in a single day, for 2 days, was evaluated. An improvement in the performance in the second day was considered motor learning.

### The three-chamber test

The three-chamber test was evaluated in a tripartite arena (Stoelting). The test was split in three epochs. In the first part, the animal was allowed to freely explore the three chambers of the arena for 20 minutes. In the second part the animal was allowed to voluntarily interact with an empty gridded recipient or with a similar recipient containing a stranger animal, for 10 minutes. The preference index for social behavior (S1:E) was determined as follows: 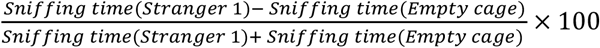. In the third part of the trial, which also lasted 10 minutes, a second stranger was placed in the previously empty recipient. The preference index for social novelty (S2:S1) was determined as follows: 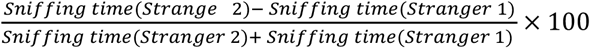. The time spent in close proximity with the gridded recipients was evaluated using the Observer XT 12 software (Noldus, The Netherlands).

### Slice preparation and electrophysiological recordings

WT and stargazin V143L KI mice littermates (P15-P20) were deeply anesthetized with isofluorane and transcardially perfused with ice-cold sucrose cutting solution (212.7 mM sucrose, 2.6 mM KCl, 1.23 mM NaH2PO4, 26 mM NaHCO3, 10 mM glucose, 3 mM MgCl2, 1mM CaCl2, pH 7.4, 300–320 mOsm) oxygenized with carbogen (95% O2 and 5% CO2). The brain was quickly removed and immersed in oxygenated ice-cold sucrose cutting solution. 300 µm acute hippocampal sagittal slices were prepared using a vibratome (Leica VT1200s, Leica Microsystems, USA). The slices were collected and transferred to a submersion holding chamber with artificial cerebrospinal fluid (aCSF) continuously oxygenated with carbogen, at 32°C, for 30 minutes. After that, slices were allowed to further recover for 1 hour at room temperature, in oxygenated aCSF, before recording. aCSF composition (mM) for whole-cell patch-clamp recordings: 125.1 NaCL, 2.5 KCl, 1.1 NaH2PO4, 25 NaHCO3, 25.0 glucose, 0.5 MgSO4, 2 CaCl2, pH 7.4, 300–310 mOsm. aCSF composition (mM) for field recordings: 130.9 NaCL, 2.5 KCl, 1.1 NaH2PO4, 24.0 NaHCO3, 12.5 glucose, 0.5 MgSO4, 2 CaCl2, pH 7.4, 300–310 mOsm.

CA1 pyramidal neurons were visualized under infrared-differential interference contrast (IR-DIC) microscopy using an upright microscope (Axio Examiner.D1, Zeiss, Germany). Whole-cell voltage-clamp recordings were performed at a holding potential of −80 mV using a Multiclamp 700B amplifier, digitized at 20 kHz with Digidata 1550A (Molecular Devices Corporation), and acquired using Clampfit 10.7 software (Axon Instruments). Slices were kept in a recording chamber perfused with oxygenated aCSF (2–3 mL/min), at 30°C, supplemented with 1 µM TTX, 100 µM picrotoxin and 50 µM D-APV, to isolate AMPAR-mediated mEPSC. Borosilicate glass recording pipettes (3-5 MΩ) were filled with a Cs-based solution (115.0 mM CsMeSO3, 20.0 mM CsCl, 2.5 mM MgCl2, 10.0 mM HEPES, 0.6 mM EGTA, 10 mM Na-phosphocreatine, 4 mM ATP sodium salt, 0.4 mM GTP sodium salt, pH 7.3, 295–300 mOsm). Data were filtered at 2 kHz. Cells were discarded if Ra was higher than 25 MΩ or if holding current or Ra changed more than 20%. Data were analysed using Clampfit software (Axon Instruments) using a template search method to detect events. Experiments and analysis were done blind to the genotype.

fEPSPs were evoked by stimulating the Schaffer collaterals at 0.05 Hz using a bipolar electrode (100 µs stimulus; Bowdoin, ME, USA) connected to a stimulator Digitimer model DS3 (Digitimer, UK) and recorded in CA1 *stratum radiatum*. Recordings were performed at 25°C in a recording chamber constantly perfused with oxygenated aCSF (2–3 mL/min). The recording pipette was filled with aCSF (2-4 MΩ). An input-output curve, starting at 20 µA with 10 µA increments, was performed and the stimulation intensity was set to elicit 40-50% of the maximal response. Only slices displaying a stable signal response over a period of 10 minutes were used. Short-term synaptic plasticity was assessed by measuring paired-pulse facilitation (PPF) using a standard protocol, as previously described ^63^. LTP was induced by theta-burst stimulation (TBS; 10 bursts of 4 stimuli at 100 Hz with a burst frequency of 5 Hz) ^64^. A baseline was recorded in the current-clamp mode with a single stimulation at 0.05 Hz (100 µs stimulus) for 15 minutes immediately before TBS. Changes in fEPSPs were recorded at 0.05 Hz for 60 minutes after TBS. Recordings were filtered at 0.1 Hz-1 kHz and digitized at 10 kHz. For each data point three individual traces were averaged. Fiber volley amplitude and synaptic response slopes were analysed using Clampfit software.

### Labelling, detection and morphological classification of dendritic spines

To achieve sparse labelling of neurons in the hippocampus, we performed tail-vein injections in 4-week-old animals, with 5 µL of AAV9.Syn.eGFP.WPRE.bGH at a titer of 8.88×10^12^ (Penn Vector Core, University of Pennsylvania, PA) diluted in sterile PBS to a final volume of 100 µL. Four weeks post-injection, animals were sacrificed and the brains were collected and processed for neuronal imaging as already described. Brains were sliced in the cryostat to obtain 100 μm serial coronal slices and the GPF fluorescence signal was enhanced by performing immunostaining against GFP. Sections were then mounted in gelatinized slides using Vectashield with DAPI (Vector Laboratories, USA). Images of secondary basal and apical dendrites from CA1 pyramidal neurons expressing GFP were acquired in an LSM 710 Confocal microscope (Zeiss, Germany) with a Plan Apochromat 63x (NA 1.4) oil objective. The dendritic segments imaged were randomly selected from at least four different sections. Per each animal, eight basal and eight apical dendritic segments, of approximately 20 µm, from different cells were imaged. The z-stack images were deconvolved using Huygens software (Scientific Volume Imaging, Netherlands) and spines were visualized and identified using Imaris software (Bitplane, Switzerland). Spines were manually categorized into five groups based on its morphology: mushroom (defined neck and a large head), stubby (without a defined neck), branched (cup-shaped; with a head protrusion; with multiple heads), thin (thin neck and small head) and filopodia (without a defined head). Image acquisition and analysis was performed by a blind-to-genotype observer.

### Electron microscopy

Sample preparation and post-synaptic density parameter measurements were performed as previously described in ^63^. Eight-week-old mice were anesthetized with isofluorane and transcardially perfused with ice-cold PBS followed by 4% paraformaldehyde. Cortices and hippocampi were dissected, and small punches of tissue were left overnight in PFA 4% and then transferred into a 2.5% glutaraldehyde solution in 0.1 M sodium cacodylate buffer (pH 7.2), where they were kept at 4°C overnight. The tissue was then rinsed in a cacodylate buffer and post-fixed with 1% osmium tetroxide for 1 h. After rinsing in buffer and distilled water, 1% aqueous uranyl-acetate was added to the tissues, in the dark, during 1 h for contrast enhancement. Following rinsing in distilled water, samples were dehydrated in a graded acetone series (70–100%) and then impregnated and included in Epoxy resin (Fluka Analytical). Ultrathin sections (70 nm) were mounted on copper grids and observations were carried out on a FEI-Tecnai G2 Spirit Bio Twin at 100kV. PSD measurements were performed using ImageJ (NIH, Bethesda, Maryland) by a blind-to-genotype observer.

### Tissue lysates and post-synaptic density isolations

Eight-week old WT, stargazin KI^+/VL^ and KI^VL/VL^ mice were anesthetized with isoflurane and euthanized by decapitation. Tissue lysates and post-synaptic density (PSD) isolations were carried out as described below. All procedures were performed at 4 °C.

### Hippocampal lysates

Hippocampi from WT, KI^+/VL^ and KI^VL/VL^ mice were mechanically homogenized in TEEN buffer (25 mM Tris pH 7.4, 1mM EDTA, 1 mM EGTA, 150 mM NaCl and 1% Triton X-100, supplemented with 1 mM DTT, 0.2 mM PMSF, 1 µg/ml CLAP (1 mg/ml of Chymostatin, Leupeptin, Antipain and Pepstatin), 5 mM NaF and 0.1mM Na3VO4), using a motor driven glass-Teflon homogenizer at 900 rpm (50 strokes). Hippocampal homogenates were centrifuged at 700 g for 10 minutes and the supernatants were collected and sonicated using an ultrasonic probe for 60 seconds (6 pulses of 5 seconds). The samples were again centrifuged at 21100 g for 10 minutes and the supernatants were collected.

### Whole brain lysates and PSD isolations

Cortices and whole brain samples from WT, KI^+/VL^ and KI^VL/VL^ were dissected and homogenized in HEPES A buffer (4 mM HEPES pH=7.4, 0.32 M sucrose, supplemented with 1 mM DTT, 0.2 mM PMSF, 1 µg/ml CLAP, 5 mM NaF and 0.1mM Na3VO4), using a motor driven glass-Teflon homogenizer at 900 rpm (50 strokes). These homogenates were centrifuged at 700 x g for 15 minutes. The whole brain lysates and a fraction of the cortical lysates were collected in 2% SDS and 2.5 M Urea and stored at −80 °C for later analysis. The remaining cortical lysate was subjected to the PSD isolation protocol previously described in ^65^. Briefly, the lysates were centrifuged at 18000 x g for 15 min, resulting in the crude synaptosomal pellet, which was re-homogenized in HEPES A buffer and further centrifuged at 25000 x g for 20 min. This yielded the lysed synaptosomal membrane pellet. A fraction of this synaptosomal membrane lysate (SML) was collected in HEPES B buffer (50 mM HEPES pH7.4, 2 mM EDTA, 0.5% Triton X-100, supplemented with 1 mM DTT, 0.2 mM PMSF, 1 µg/ml CLAP, 5 mM NaF and 0.1mM Na3VO4) and 2% SDS and stored at −80 °C for later analysis. The remaining synaptosomal membrane fraction was resuspended and incubated in HEPES B buffer for 15 min and centrifuged at 32000 x g for 20 min. The resulting pellet was resuspended and incubated in HEPES B buffer for 15min and centrifuged at 200000 x g for 20 min. The resulting pellet, consisting of the PSD fraction, was collected in HEPES B buffer and 2% SDS, 2.5 M urea and stored at −80 °C for later analysis.

### Cortical lysates for immunoprecipitation

Cortical lysates for immunoprecipitation were obtained as previously described ^66^. Briefly, fresh cortices from WT or KI^VL/VL^ littermates were mechanically homogenized in 5 ml of buffer A (20 mM HEPES, 0.15 mM EDTA, 0.4 mM EGTA, 10 mM KCl, pH 7.5, supplemented with1 mM DTT, 0,2 mM PMSF, 1 µg/ml CLAP, 5 mM NaF, 0, 1mM Na3VO4) followed by sonication and then centrifuged for 10 min at 860 × g. The resulting supernatant was centrifuged for 30 min at 17000 × g. Buffer A was supplemented with 15% sucrose and 5 ml was used to homogenize each pellet with 20 strokes, which was further centrifuged for 10 min at 860 × g to remove genomic DNA. The brain membranes present in the supernatant were centrifuged again for 30 min at 17000 × g. The pellets were solubilized in buffer B (20 mM HEPES, 1% Triton-X100, 150 mM NaCl, 0.15 mM EDTA, 4 mM EGTA, pH 7.5, with the same cocktail of protease and phosphatase inhibitors) with 20 dunces with a potter and centrifuged at 17000 x g for 45 minutes, yielding the cortical lysates.

### Stargazin immunoprecipitation

Immunoprecipitation (IP) of stargazin was performed as previously described ^66^. Briefly, protein concentration of cortical lysates was quantified with a Bicinchoninic Acid (BCA) assay (Fisher Scientific, USA). The lysates (500 µg) were incubated with the anti-stargazin antibody (IP+) (AB-9876, Merck Millipore, 2 µg) or with normal Rabbit polyclonal IgG (IP-) (12-370, Merck Millipore, 2 µg) for 1 h at 4 °C under rotation and then incubated overnight with 50 µl of protein-A Sepharose at 4 °C. Resin was washed with 1 ml of buffer B and 0,5 ml of the same buffer supplemented with 500 mM NaCl. Beads were resuspended in 50 µl of 2x denaturing buffer (62.5 mM Tris·HCl (pH 6.8), 10% Glycerol, 2% SDS, 0.01% bromophenol blue, and 5% β-mercaptoethanol).

### Lambda phosphatase treatment

Lambda phosphatase (λ-PP) treatment of cortical PSD samples was performed using the λ-PP treatment kit from New England Biolabs (USA), according to the manufacturer’s instructions. In brief, cortical lysates from WT, stargazin KI^+/VL^ and KI^VL/VL^ mice were obtained according to the previously described protocol, without the supplementation with the phosphatase inhibitors NaF and Na3VO4. The lysates were then divided into two groups: treated and untreated samples. The cortical lysates from the untreated group were supplemented with 5 mM NaF and 0.1mM Na3VO4. Samples from the treated group were supplemented with 12.5% of NEBuffer for Protein MetalloPhosphatases (PMP), 12.5% of 10mM MnCl2 and 2.5% of Lambda Protein Phosphatase. All samples from both groups were then incubated at 30 °C for 30 min and processed according to the previously described PSD isolation protocol.

### SDS-PAGE and Western blot

Protein quantification was performed using the BCA assay (Fisher Scientific, USA). Samples were denatured with sample buffer 5x (NZYTech, Portugal) and resolved by SDS-PAGE in Tris-glycine-SDS buffer (25 mM Tris, 192 mM glycine, 0.1 % SDS, pH 8.3) in an 11% polyacrylamide gel. Stargazin immunoprecipitation samples were resolved in 4-20% Mini-PROTEAN® TGX™ Precast Protein Gels (BioRad) in the same Tris-glycine-SDS buffer. All SDS-PAGE gels were subjected to an overnight electrotransfer (40 V, 4 °C) to a PVDF membrane (Millipore, USA). The membranes were then blocked using a 5% milk solution in TBS (20 mM Tris, 137 mM NaCl, pH 7.6) supplemented with 0.1% Tween-20 (TBS-T) for 1h at room temperature (RT). After blocking, the membranes were incubated with the primary antibodies against stargazin (AB-9876, Merck Millipore, 1:750 in 3% Milk TBS-T), GluA1 (MAB2263, Merck Millipore, 1:1000 in 5% Milk TBS-T), GluA2 (MAB397, Merck Millipore, 1:1000 in 5% Milk TBS-T) and PSD95 (MA1-045, ThermoFisher Scientific,1:1000) for 2h at RT. The membranes were washed 3 times for 10 min in TBS-T and then incubated with the appropriate alkaline phosphatase-conjugated secondary antibody (#115-055-146 or #211-055-109, Jackson ImmunoResearch, 1:10000 in 5% milk TBS-T) for 45 min at RT. Following 3 washes in TBS-T, membranes were developed with the alkaline phosphatase substrate ECF (GE Healthcare, USA) and the fluorescent signal was acquired using a ChemiDoc Gel Imaging System (Bio-Rad, USA). The results were analyzed using ImageJ (NIH, Bethesda, Maryland).

### Statistical analysis

The normality of population distributions was calculated for each experiment by comparison with a theoretical normal distribution using the Shapiro-Wilk normality test. According to this evaluation parametric or non-parametric tests were used, as described in the figure legends. For all tests, *p* < 0.05 was considered statistically significant. Analyses were performed using GraphPad (Prism).

## Supporting information

Caldeira_Suppinfo

## ACKNOWLEDGEMENTS

This work was supported by a NARSAD Independent Investigator Grant (#23151) and a NARSAD Young Investigator Grant (#20733) from the Brain and Behavior Research Foundation, by a research grant from the Jérôme Lejeune Foundation (#1530), by a Marie Curie Integration Grant (618525), by a Bial Foundation Grant (266/2016), by national funds through the Portuguese Science and Technology Foundation (FCT: UID/NEU/04539/2013, UIDB/04539/2020, POCI-01-0145-FEDER-28541, POCI-01-0145-FEDER-016682, PTDC/QUI-OUT/32243/2017 and CPCA/A0/7302/2020), and by the European Regional Development Fund (ERDF), through the Centro 2020 Regional Operational Programme, under project CENTRO-01-0145-FEDER-000008:BrainHealth 2020. GLC, NB, MVR, ME and CAVB were supported by FCT through Ph.D. scholarships SFRH/BD/51962/2012, SFRH/BD/144881/2019, SFRH/BD/129236/2017, SFRH/BD/51958/2012 and SFRH/BD/145457/2019, respectively. ASI and JG were supported by FCT through Post-doctoral fellowship SFRH/BPD122299/2016 and SFRH/BPD/120611/2016, respectively. RPG and RM received support from FCT/DGES, under the program “Verão com Ciência”. We thank Luisa Cortes and the MICC team for assistance with microscopy imaging; Jorge Valero and Jeannette Schmidt for designing and optimizing quantification macros in Fiji ®; Lara Franco, Nuno Fonseca and Orsolya Antal for technical help. R1 ES cells for mice generation were a kind gift from Dr. Andras Nagy (Mount Sinai Hospital); Stargazin plasmids were a kind gift from Dr. Daniel Choquet (IINS, Bordeaux). Schematic figures were created using Biorender.com. We thank all Ana Luísa Carvalho’s lab members for technical assistance and for the indispensable discussion of the work.

## AUTHOR CONTRIBUTIONS

Conceptualization, ALC, JP, ISM, GLC, ASI; Methodology, ALC, JP, GLC; Investigation, GLC, ASI, NB, CAVB, MVR, TR, RM, RPG, BC, SRL; Writing – Original Draft, ALC, GLC, ASI, NB, ISM; Funding Acquisition, ALC, JP, ISM; Supervision, ALC, JP, ISM.

