## Supplementary material for "Aberrant hippocampal transmission and behavior in mice with a stargazin mutation linked to intellectual disability": Caldeira_Suppinfo

### SUPPLEMENTARY METHODS

#### RNA extraction and qPCR

Total RNA was extracted from 8 weeks-old mice cortices and hippocampi with miRCURY™ RNA Isolation Kits - Cell & Plant (Exiqon), according to manufacturer's instructions. RNA concentration was determined using a NanoDrop 2000c/2000 UV-Vis Spectrophotomer. cDNA was synthesized by reverse transcription NZY First-Strand cDNA Synthesis Kit (NZYtech), according to manufacturer instructions, in T100 thermal cycler (Bio-Rad).

Primers for real-time PCR were designed using the Universal Probe library (Roche Life Science). Gene expression analysis was performed using SsoFast SuperMix (Bio-Rad), according to manufacturer's specifications. The fluorescence was measured using the iQ5 Multicolor Real-Time PCR Detection System (BioRad). Data was analysed with the GenEx software (MultiD Analyses, Sweden). The constitutively expressed housekeeping gene encoding HPRT was used as control.

### Behavior analyses

#### *Open field test*

In the open field test animals were allowed to freely move, for one hour, in an opaque arena (40x40 cm). The room was illuminated by white LED lamps at 100 lux. The locomotor activity (total distance traveled, instant velocity and time in center zone) was quantified with the Bonsai software using the recorded videos.

#### *Elevated-plus maze test*

The EPM consists of a cross-shaped platform, containing four arms, which is elevated from the ground; two of the arms are enclosed by walls whereas the other two are open and thus subjected to bright illumination (100 lux). The animals initiated the test in the center of the maze and their movement was recorded for 10 minutes. The preference for the open or closed arms, as well as the time spent in the center was evaluated using the Bonsai software.

#### *Forced-swimming test*

FST was conducted by placing the mice in 2 L glass beakers filled with 1.6 L of water at 24 ± 1°C. The total duration of immobility over the 6 minutes observation period was scored using the Observer XT 12 software (Noldus, Netherlands). Immobility was defined as the lack of motion of the whole body, except for small movements necessary to keep the animal's head above the water.

#### *T-maze spontaneous alternation test*

The spontaneous alternation test was performed in a T-shaped maze (30x10 cm), illuminated with 15-20 lux. The mice were introduced in the start arm and allowed to choose one of the other arms. After the animals entered one of the arms, it was enclosed in that area with a sliding door, for 30 seconds. The animal was then removed from the maze for 30 seconds more and re-placed in the start arm. If the animals chose the previously unexplored arm, spontaneous alternation was considered. A total of five trials were conducted in 2 consecutive days.

***Nest-building test***

Mice were transferred to individual standard home cages (20×26×13 cm) with corn bedding, one hour before the dark phase. A cotton disk was added to the cage and, 16 hours later, the nests were evaluated by blind-to-genotype observers, using a 5-point rating [1].

**3D neuronal reconstruction and Sholl analysis**

CA1 pyramidal neurons expressing GFP (see the section Labelling, detection and morphological classification of dendritic spines in the main materials and methods) were randomly selected and imaged in an LSM 710 Confocal microscope (Zeiss, Germany) using a Plan-Apochromat 20x air objective (NA 0.8). Six neurons selected from at least four different sections were analysed per animal. 3D neuronal reconstructions and morphometric analysis were performed using Imaris software (Bitplane, Switzerland). Sholl analysis was performed by quantifying the number of intersections between dendrites and concentric spheres with a radius increment of 10 μm from the cell soma. The total length of basal and apical dendrites was also determined. Image acquisition and analysis was performed by a blind-to-genotype observer.

### SUPPLEMENTARY FIGURES

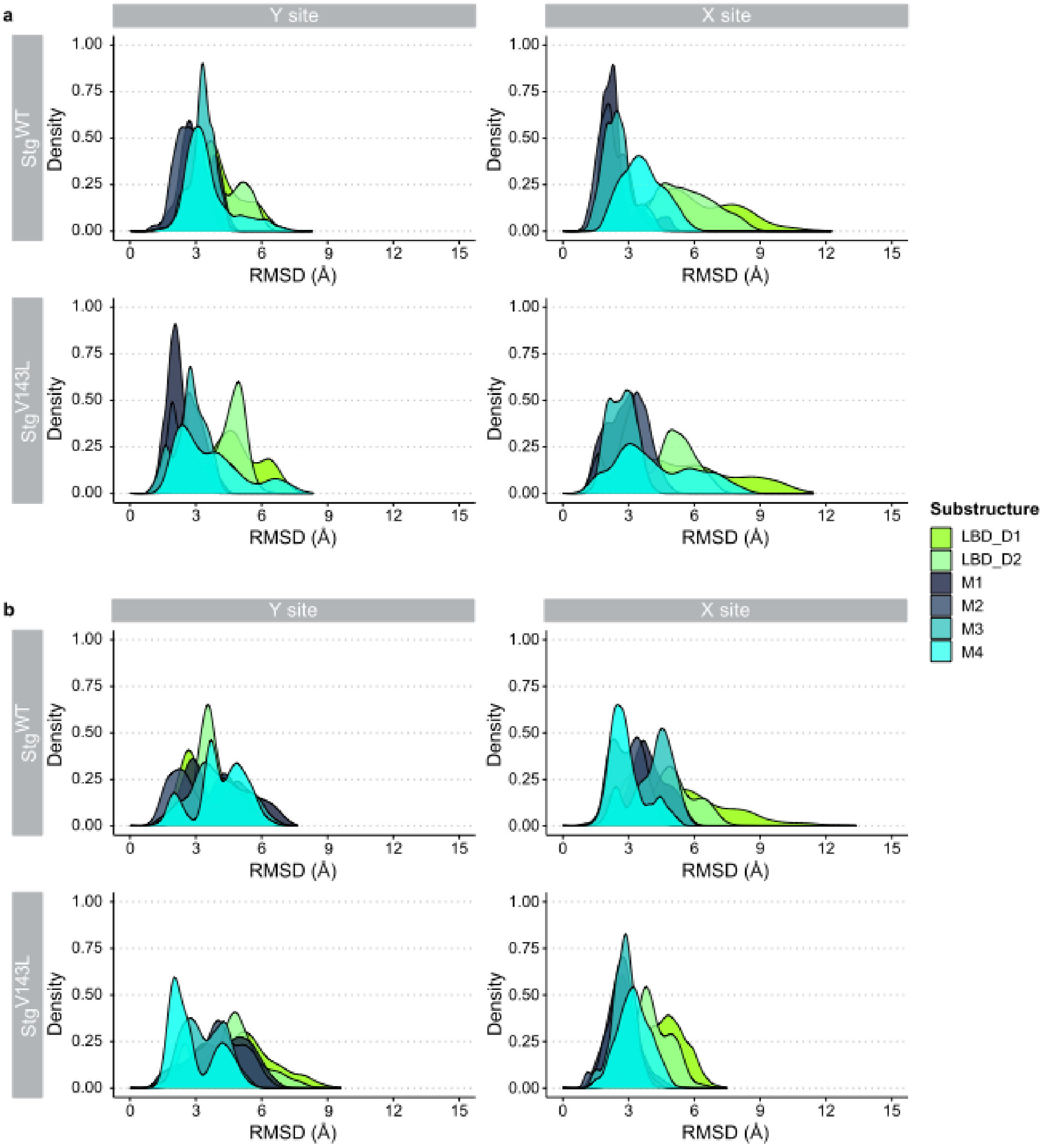


**Figure S1. RMSD density plots for Cα atoms of *Main GluA2* and *Auxiliary GluA2*.**

RMSD values were calculated for every substructure of *Main GluA2* (**a**) and *Auxiliary GluA2* (**b**) for MD simulations of both stargazin WT- and stargazin V143L-containing systems. Colors from Figure 1 were used for substructures of GluA2.

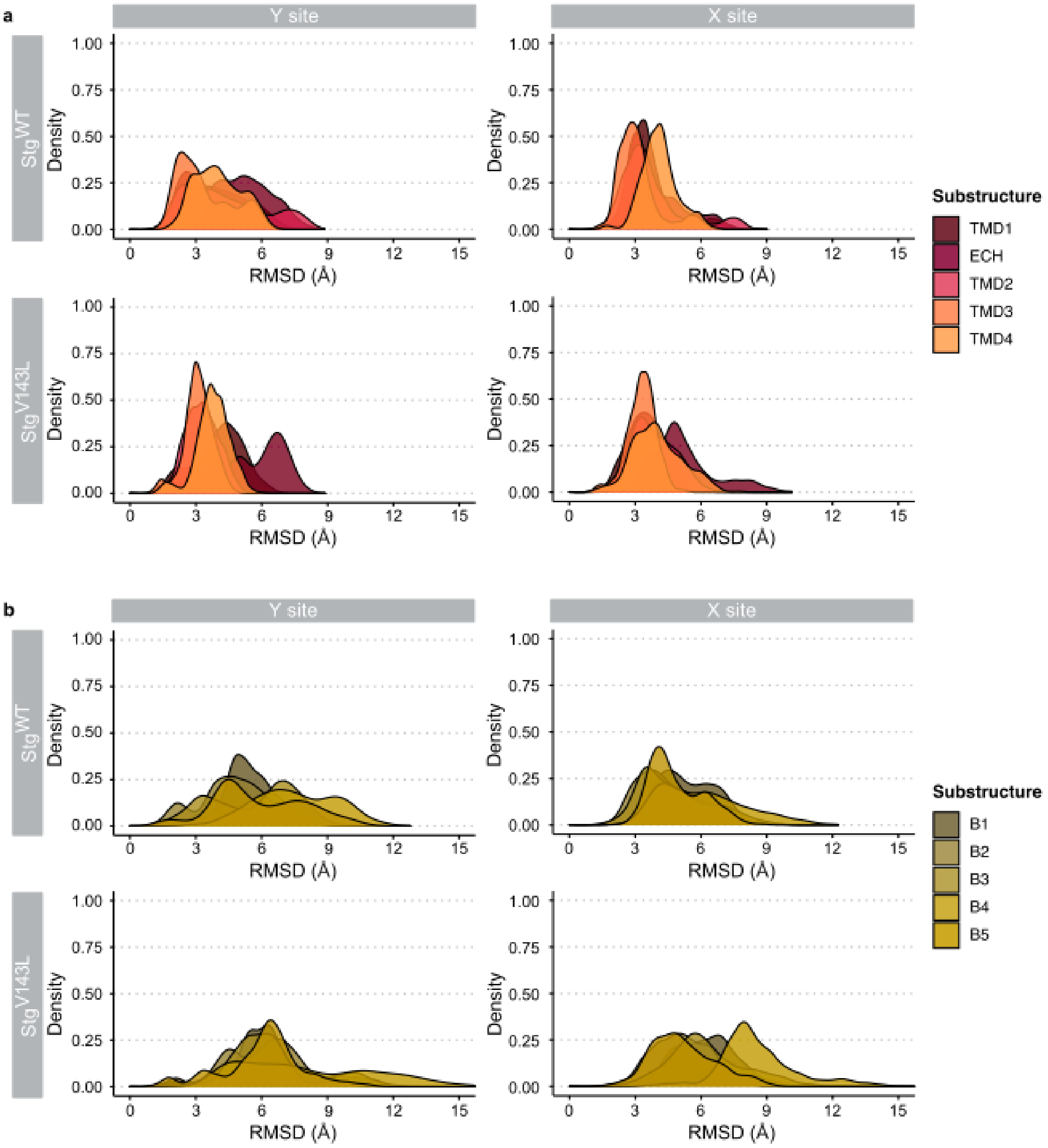


**Figure S2. RMSD density plots for Cα atoms of stargazin helices and stargazin β-strands.**

RMSD values were calculated for every helix of stargazin (**a**) and β-strands (**b**) for MD simulations of both stargazin WT and stargazin V143L-containing systems. Colors from Figure 1 were used for substructures of stargazin.

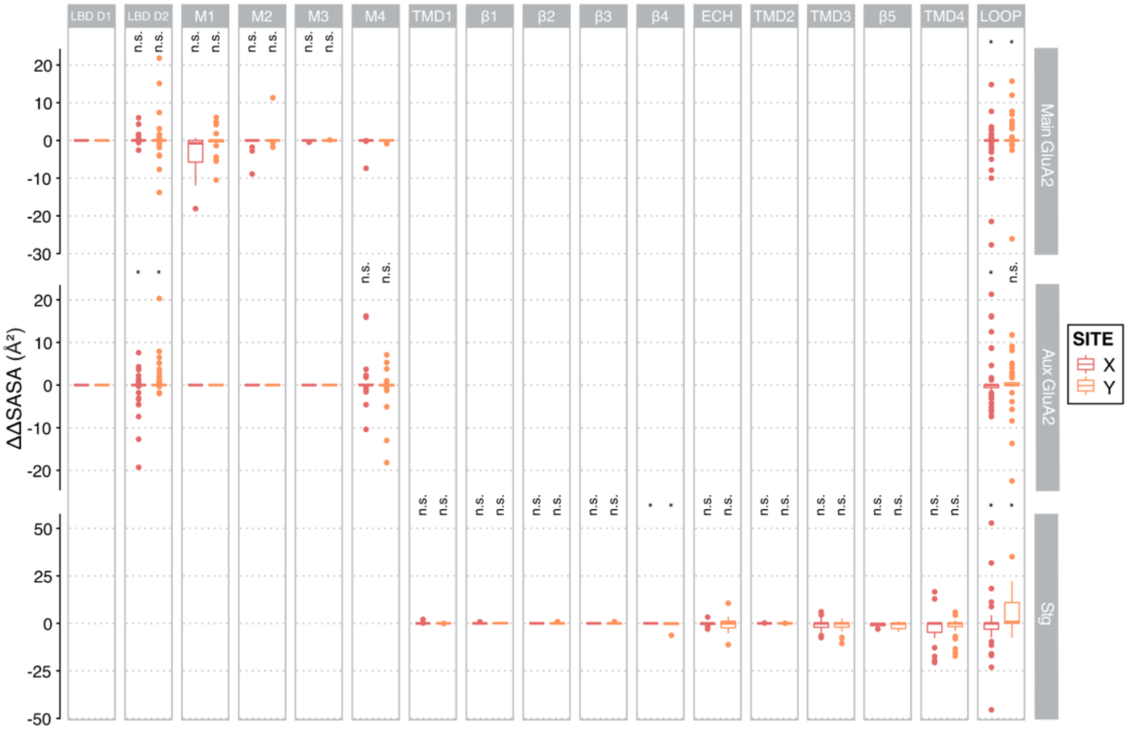


**Figure S3. ΔΔSASA values for AMPAR:stargazin complex per substructure.**

Mean ΔΔSASA of each residue was calculated as $\Delta\Delta SASA = \Delta SASA_{V143L} - \Delta SASA_{WT}$. Residues were grouped by substructure. Wilcoxon rank signed test was used to calculate statistical significance, $*p < 0.05$. X site – red; Y site – orange.

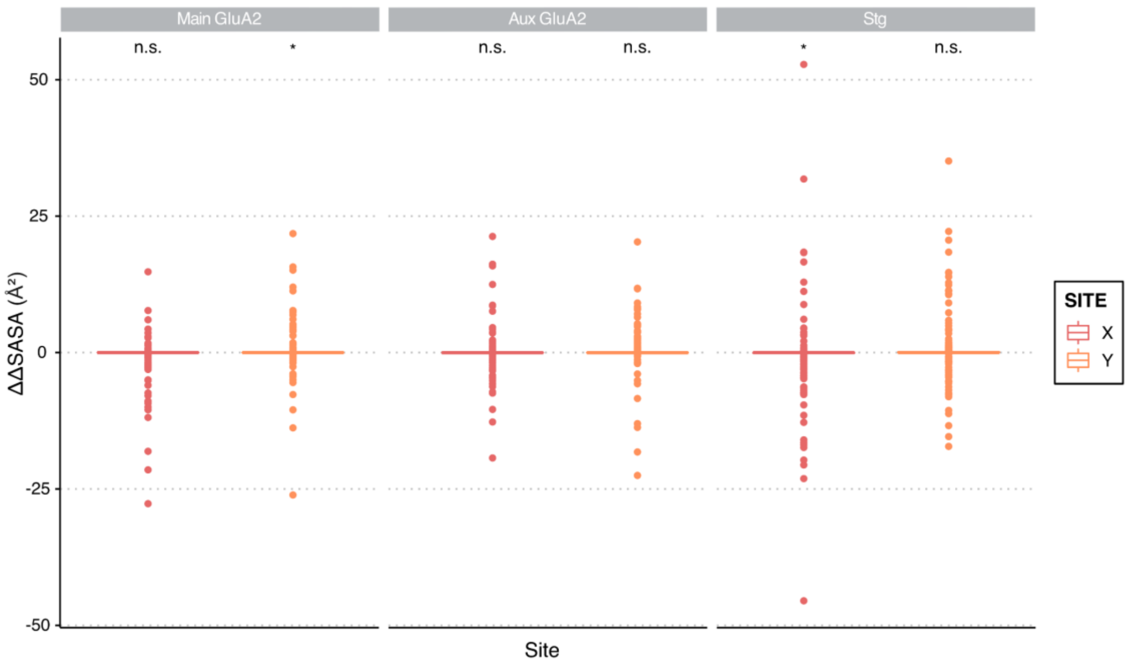


**Figure S4. ΔΔSASA values for AMPAR:stargazin complex per structure.**

Mean ΔΔSASA of each residue was calculated as $\Delta\Delta SASA = \Delta SASA_{V143L} - \Delta SASA_{WT}$. Residues were grouped by structure. Wilcoxon rank signed test was used to calculate statistical significance, *$p < 0.05$. X site – red; Y site – orange.

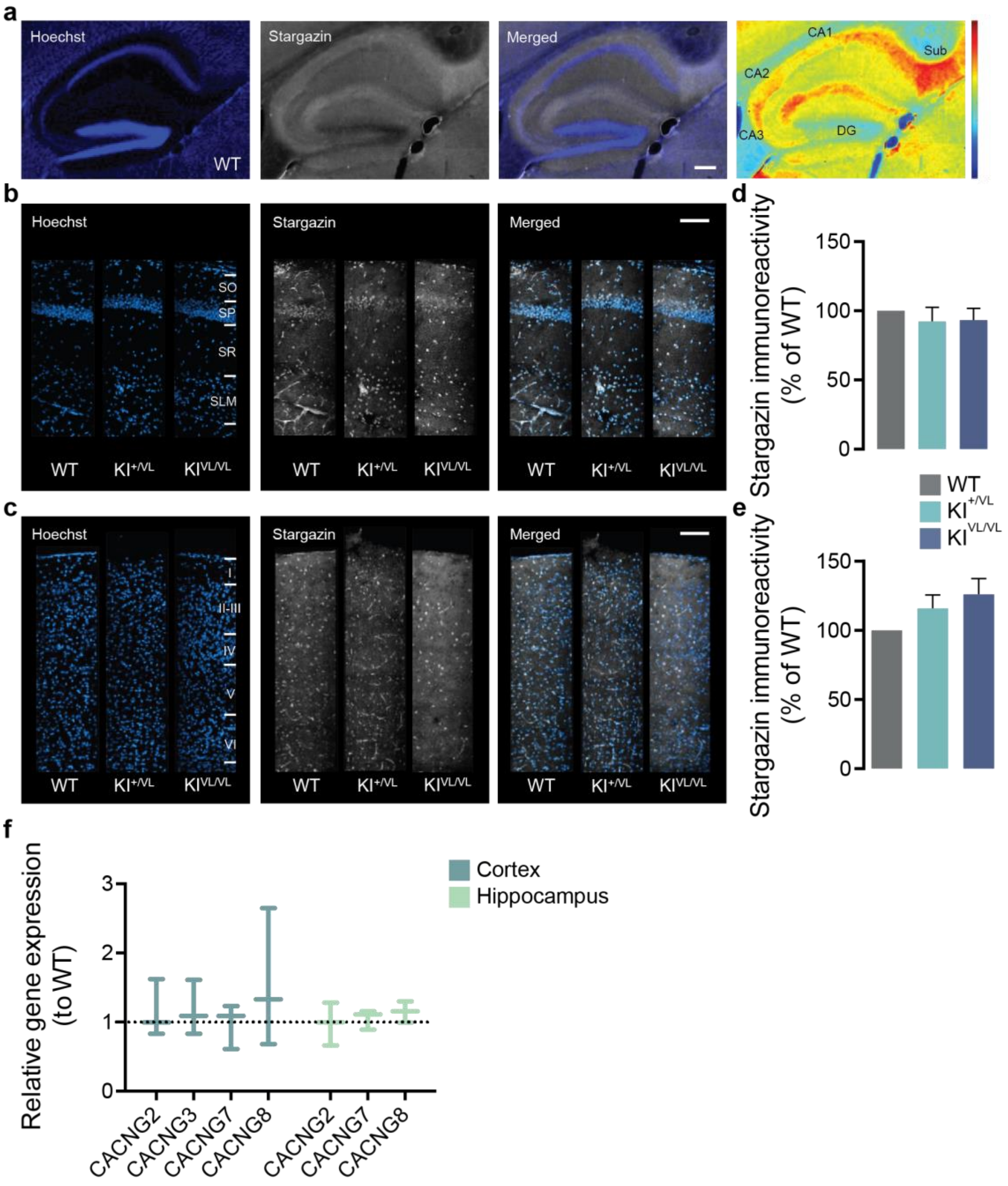


**Fig. S5. Stargazin expression is not affected by the V143L mutation.**

**(a)** Immunostaining of a sagittal section of a 2-month-old WT mouse and pseudo-colored image of stargazin distribution in the hippocampus. Hot colors indicate higher protein expression levels. Within the hippocampus stargazin is particularly localized in the *stratum lacunosum moleculare* (SLM) and *stratum oriens* (SO) of the CA1 and CA2 regions and more so in the *subiculum* (Sub). Nuclei were stained with Hoechst 33342. Scale bar represents 200 µm. DG, dentate gyrus.

(**b,c**) Representative confocal images of stargazin immunostaining in sagittal slices of 2-month-old WT and stargazin V143L KI mice showing a similar expression pattern in the hippocampal CA1 region (**b**) and in the cortex (**c**) of stargazin V143L animals compared to WT controls. No alterations in lamination of the cortex and hippocampus were observed. Nuclei were stained with Hoechst 33342. Scale bar represents 100 µm. SO, *stratum oriens*; SP, *stratum pyramidale*; SR, *stratum radiatum*; SLM, *stratum lacunosum moleculare*. (**d,e**) Immunostaining of stargazin by Western blot (see Fig. 3e) showed no significant alterations of its levels in hippocampal (**d**) and cortical (**e**) lysates from stargazin V143L KI mice compared to WT controls. Data are presented as mean ± SEM. One-sample t-test to the value of 100%. n = 6 for hippocampal samples and n = 9 for cortical samples for all genotypes. (**f**) Relative gene expression of *Cacng2*, *Cacng3*, Cacng7 and *Cacng8* is not altered in the cortex of stragazin $KI^{VL/VL}$ animals, in comparison to WT animals. The expression of *Cacng2*, *Cacng7* and *Cacng8* was not altered in the hippocampus of $KI^{VL/VL}$ animals, in comparison to WT. Gene expression levels of $KI^{VL/VL}$ animals are normalized for the expression levels in WT littermates. The expression level of each gene was normalized to the expression of the control gene *Hprt* in the corresponding condition. Data are presented as median and range. n=3 for all conditions.

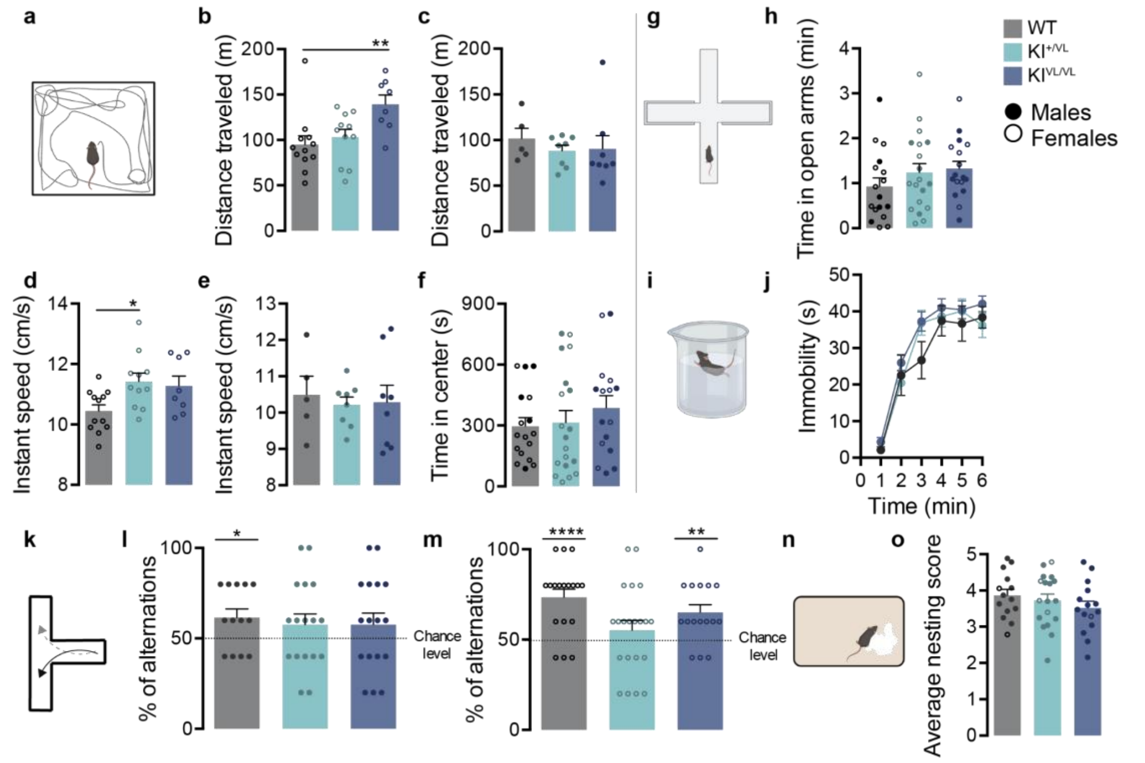


**Figure S6. Stargazin V143L KI mice present hyperactivity and working memory deficits.**

(**a**) The locomotor activity was evaluated using the open field test. (**b**) Female stargazin $KI^{VL/VL}$ animals travelled significantly more than WT or heterozygous animals. (**c**) No changes were observed among male animals from different genotypes. (**d**) The instant speed of stargazin $KI^{+/VL}$ female mice was significantly higher than that of WT female mice, but (**e**) did not differ among males from the three genotypes. (**f**) Mutant animals did not present changes in the time spent in the center of the open field arena, (**g,h**) in the open arms of the elevated plus maze, or (**i,j**) in the immobility time in the forced swimming test. Together, these data suggest that these animals do not present anxiety-like behaviors. Data are presented as mean ± SEM.*p < 0.05, **p <0.01. Kruskal-Wallis test, n ≥ 8 for females from all genotypes and ≥ 5 for males from all genotypes for open field test, n≥17 for all genotypes (males and females) for elevated plus maze test and n ≥ 8 for all genotypes (males and females) for forced swimming test. (**k**) Spontaneous alternation was evaluated using the T-maze. (**l,m**) WT animals alternated significantly more than 50% of times, whereas male stargazin $KI^{+/VL}$ and stargazin KIVL/VL animals failed to do so. Data are presented as median with range. **p < 0.01, ****p < 0.0001, One-sample Wilcoxon signed rank test against a value of 50%. n ≥ 31 (males and females) for all genotypes. (**n,o**) In the nesting behavior test, scores given by blind-to-genotype observers did not significantly vary among genotypes. One-way ANOVA followed by Dunnet's multiple comparison test, n ≥ 14 for all genotypes (males and females).

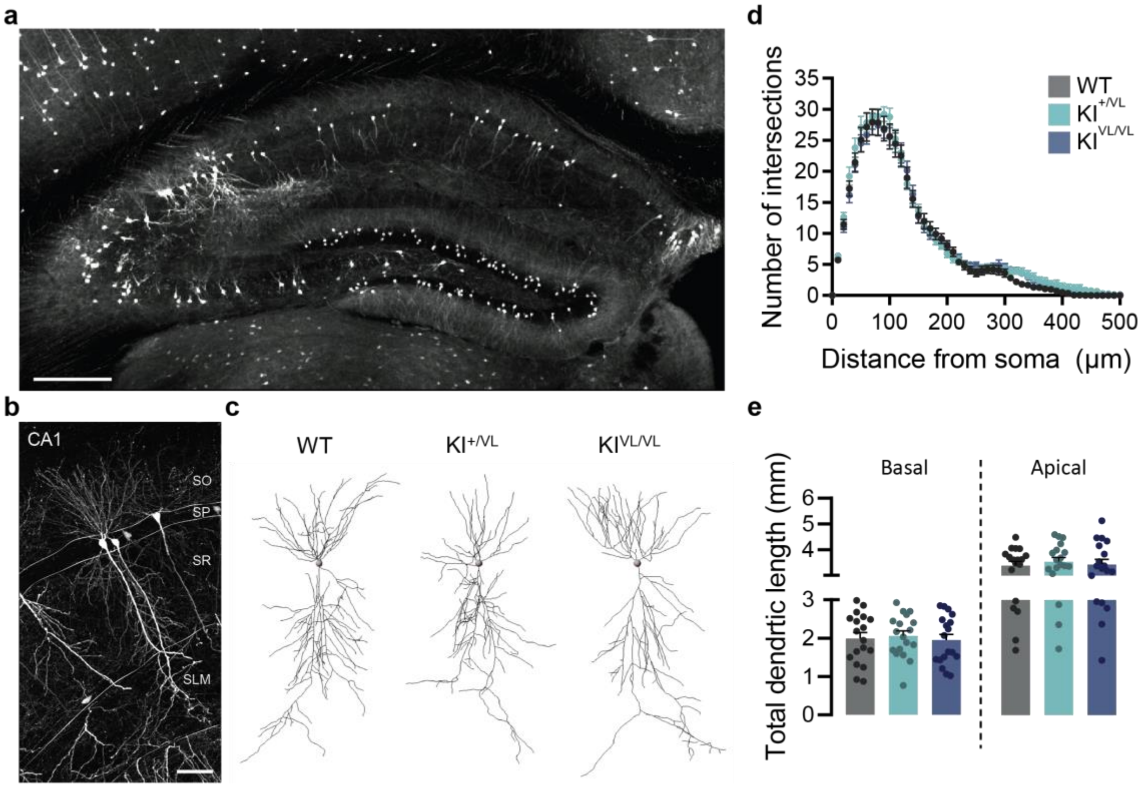


**Figure S7. The morphology of hippocampal CA1 pyramidal neurons is not altered in stargazin V143L KI mice.**

(**a**) Representative image of sparsely labelled hippocampal neurons after tail vein injection with AAV9.hSyn.eGFP. Scale bar represents 300 µm. (**b**) High-magnification representative image showing GFP-labelled neurons in the CA1 region. Scale bar represents 50 µm. (**c**) Representative images of 3D reconstructions of CA1 pyramidal neurons from WT, stargazin KI$^{+/VL}$ and stargazin KI$^{VL/VL}$ mice. (**d**) Sholl analysis showed that there are no significant changes in the dendritic tree architecture of CA1 pyramidal neurons of stargazin V143L mice. (**e**) The total dendritic length of basal and apical dendrites is similar for all genotypes. Data are presented as mean ± SEM. n = 18 neurons/3 animals for all genotypes.

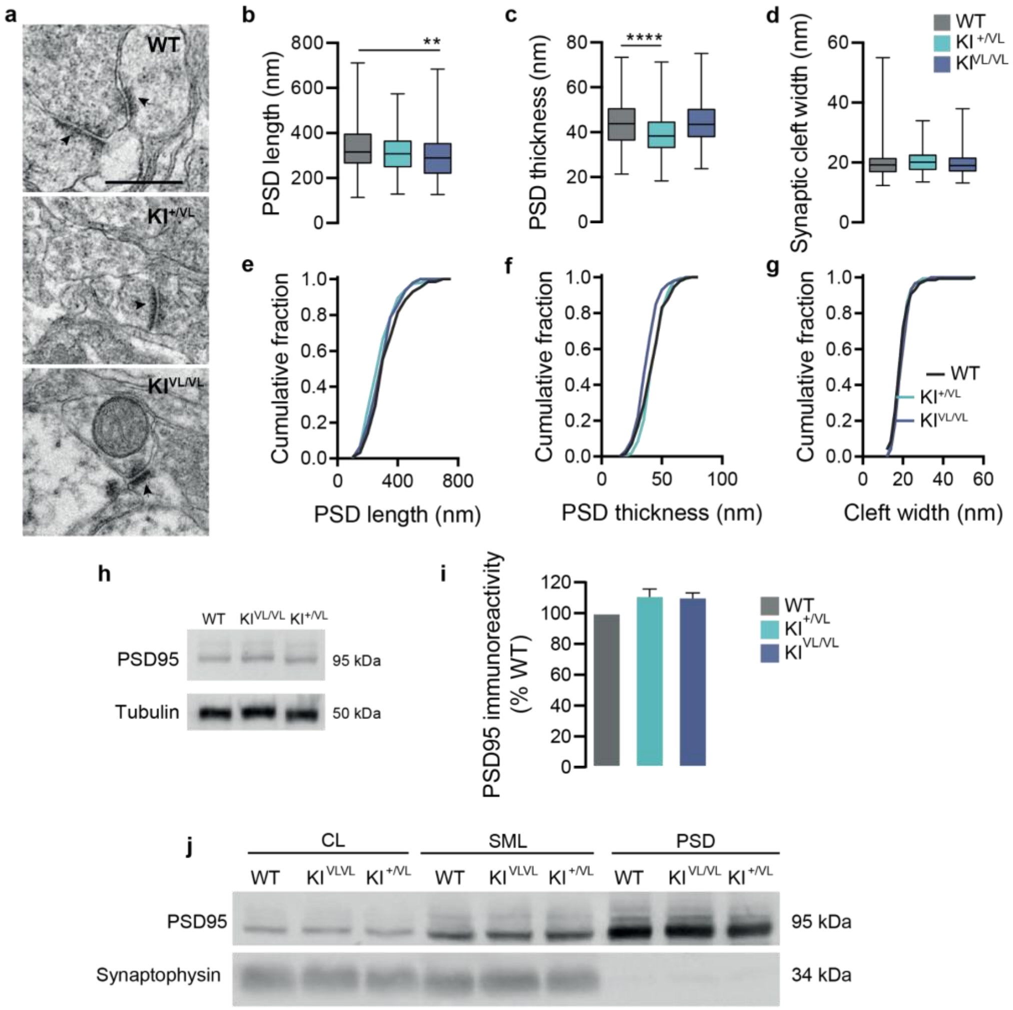


**Figure S8. Stargazin V143L KI mice display alterations in cortical post-synaptic densities ultrastructure.**

(**a**) Representative electron transmission microscopy images of cortical synapses from WT, stargazin $KI^{+/VL}$ and $KI^{VL/VL}$ animals and quantification of post-synaptic density length (**b**, **e**) and thickness (**c**, **f**) and cleft width (**d**, **g**). Data are presented as median with range. $^{**}p < 0.01$, $^{****}p < 0.0001$, Kruskal-Wallis followed by Dunn's multiple comparison post-hoc test. n = 130 PSDs/2 animals for WT mice, n = 154 PSDs/2 animals for $KI^{+/VL}$ mice, n = 152 cells/2 animals for $KI^{VL/VL}$ mice. The arrows indicate the post-synaptic densities. Scale bar represents 500 nm. (**h**) Immunostaining of PSD95 by Western blot showed no significant alterations in its levels in cortical samples from stargazin V143L

KI mice compared to WT controls. One-sample t-test to the value of 100%. (**i**) Western blot analysis of cellular lysate (CL), synaptic membrane lysate (SML) and post-synaptic densities (PSD) isolated from the brain cortex shows enrichment of PSD95 and absence of synaptophysin labeling in PSDs, confirming efficient isolation of this fraction.
